# Metabolomics of *in vivo* inflammation identifies soluble sialic acid as a conserved myeloid-cell metabolite

**DOI:** 10.64898/2026.07.06.736822

**Authors:** Helen A. Jordan, Joseph A. Tandurella, Vineeth Vengayil, Tanay S. Parnaik, Swechha Mainali Pokharel, Kathleen Mulka, Sara Cherry, E. John Wherry, Caroline R. Bartman

**Affiliations:** Department of Systems Pharmacology and Translational Therapeutics, Perelman School of Medicine, University of Pennsylvania, Philadelphia, PA, USA; Department of Pathology and Laboratory Medicine, Perelman School of Medicine, University of Pennsylvania, Philadelphia, PA, USA; Department of Pathobiology, School of Veterinary Medicine, University of Pennsylvania, Philadelphia, PA, USA; Institute for Immunology and Immune Health, University of Pennsylvania Perelman School of Medicine, Philadelphia, PA, USA; Parker Institute for Cancer Immunotherapy, University of Pennsylvania Perelman School of Medicine, Philadelphia, PA, USA; Colton Center for Autoimmunity, University of Pennsylvania Perelman School of Medicine, Philadelphia, PA, USA

## Abstract

During an immune response, metabolism changes dramatically. Metabolites are oxidized to power immune cell functions, serve as building blocks for proliferation, and act as effectors to regulate pathogen or host cells. Though metabolic changes in cultured cells have been studied extensively, metabolism changes *in vivo* are less understood. Here, we measured metabolomic changes across six mouse tissues in three models of immune activation: CpG-DNA cytokine storm, lymphocytic choriomeningitis virus infection, and polyI:C viral mimetic injection; and carried out metabolomics in cultured macrophages activated with different stimuli. We found most metabolomic changes were exclusive to either inflamed tissues or cultured macrophages, although itaconate was strongly induced in both contexts. We then mechanistically dissected the role of the soluble sialic acid N-glycolylneuraminic acid, which is highly induced in inflamed tissues yet only modestly in cultured macrophages. This metabolite rises in tissues in different models of inflammation, and the analogous human metabolite, N-acetylneuraminic acid, is increased in human patients experiencing inflammation. We found that N-glycolylneuraminic acid is produced in CD11b+ myeloid cells by cleavage of protein-bound sialic acid. However, blocking its production did not affect CpG-DNA liver inflammation or LCMV infection in mice. Therefore, these experiments identify soluble sialic acid as a conserved biomarker of inflammation in mice and humans and highlight the differences in metabolism between *in vitro* and *in vivo* models of inflammation.

## Introduction

An immune response in the body starts when pathogens or cellular damage trigger receptors on immune and tissue stromal cells^1^. These signals start an amplifying cascade that leads to proliferation and activation of immune cells including macrophages, neutrophils, and T cells. These immune cells can migrate through a tissue, engulf dying cells, directly kill pathogens, kill infected or damaged host cells, and produce inflammatory mediators including cytokine proteins and reactive oxygen species^1^.

Metabolic changes underlie the immune response. Cells consume nutrients to make ATP, then use ATP to produce proteins, proliferate, and migrate; metabolites like itaconate and reactive oxygen species serve as direct effectors of the immune response; and sickness behaviors suppress feeding and movement and thereby alter whole-body metabolism^2–7^.

While metabolic changes caused by activation of immune cells, like T cells^2,5,6,8^ or macrophages^9–14^, have been carefully studied in tissue culture, how metabolite levels change in immune responses *in vivo* is less well characterized. For example, when a mouse or human mounts an immune response, do the same metabolites change as when immune cells are stimulated in tissue culture? Does a given inflammatory stimulus induce specific metabolite changes, or do all immune stimuli converge on similar metabolite changes? Further, is there a functional role for the metabolites changed in inflamed tissues?

Here, we set out to catalog metabolite changes in the blood, liver, spleen, lung, kidney and brain in response to different inflammatory stimuli in mice. We then measured metabolomic changes in cultured stimulated macrophages, to allow direct comparison of *in vivo* and *in vitro* immunometabolic changes. We found that in mice, most metabolite changes were unique to particular inflammatory stimuli and were also not found in stimulated macrophages in culture. In contrast, stimulated macrophages changed a shared set of metabolites no matter the stimulus. We identified one previously-uninvestigated immunometabolite, the soluble sialic acid N-glycolylneuraminic acid, that increased dramatically during *in vivo* inflammation yet only slightly in stimulated macrophages. We then characterized this metabolite’s production pathway, cell type of origin, and function in inflammation. We found that N-glycolylneuraminic acid is produced by cleavage from sialylated proteins by myeloid cells, yet blocking its production did not change the immune response or pathology in inflammation. Altogether, these experiments generate a resource of metabolomic changes in inflammation *in vivo* and *in vitro* and identify soluble sialic acid as a biomarker of inflammation in mice and humans.

## Results

### A tissue inflammation metabolomic atlas

Metabolite levels change during an immune response. Immune cell proliferation, migration, cytotoxicity, and production of protein effectors requires increases in metabolic fluxes including glycolysis, the TCA cycle, and nucleotide synthesis^2,6,12,15^. In addition, certain metabolites are produced by activated immune cells as direct effectors. Key examples include itaconate^16–20^ and nitric oxide^21,22^: these metabolites are both produced by myeloid cells and exert antimicrobial effects as well as regulatory effects on host cells. However, in most cases such functional immunometabolites were identified using cultured immune cells; careful studies of metabolomic changes during immune responses in animals may reveal immune-induced metabolic changes that do not occur in tissue culture.

Here, we set out to identify how immune challenges in mice change tissue metabolite levels and whether altered metabolites influence the outcome of the immune response. First, we gave three different immune stimuli to cohorts of C57BL/6 mice (**Figure 1A-C**): a model of cytokine storm induced by repeated injection of CpG DNA^23,24^, viral infection with the chronic Clone 13 strain of the mouse virus lymphocytic choriomeningitis virus (LCMV)^25^, and injection of the viral RNA mimetic polyinosinic-polycytidylic acid (PolyI:C)^26^. We then used high-resolution liquid-chromatography mass-spectrometry (LC-MS) metabolomics to identify metabolic changes, detecting and annotating around 700 features per dataset based on mass-to-charge ratio and retention time.

**Figure 1:**
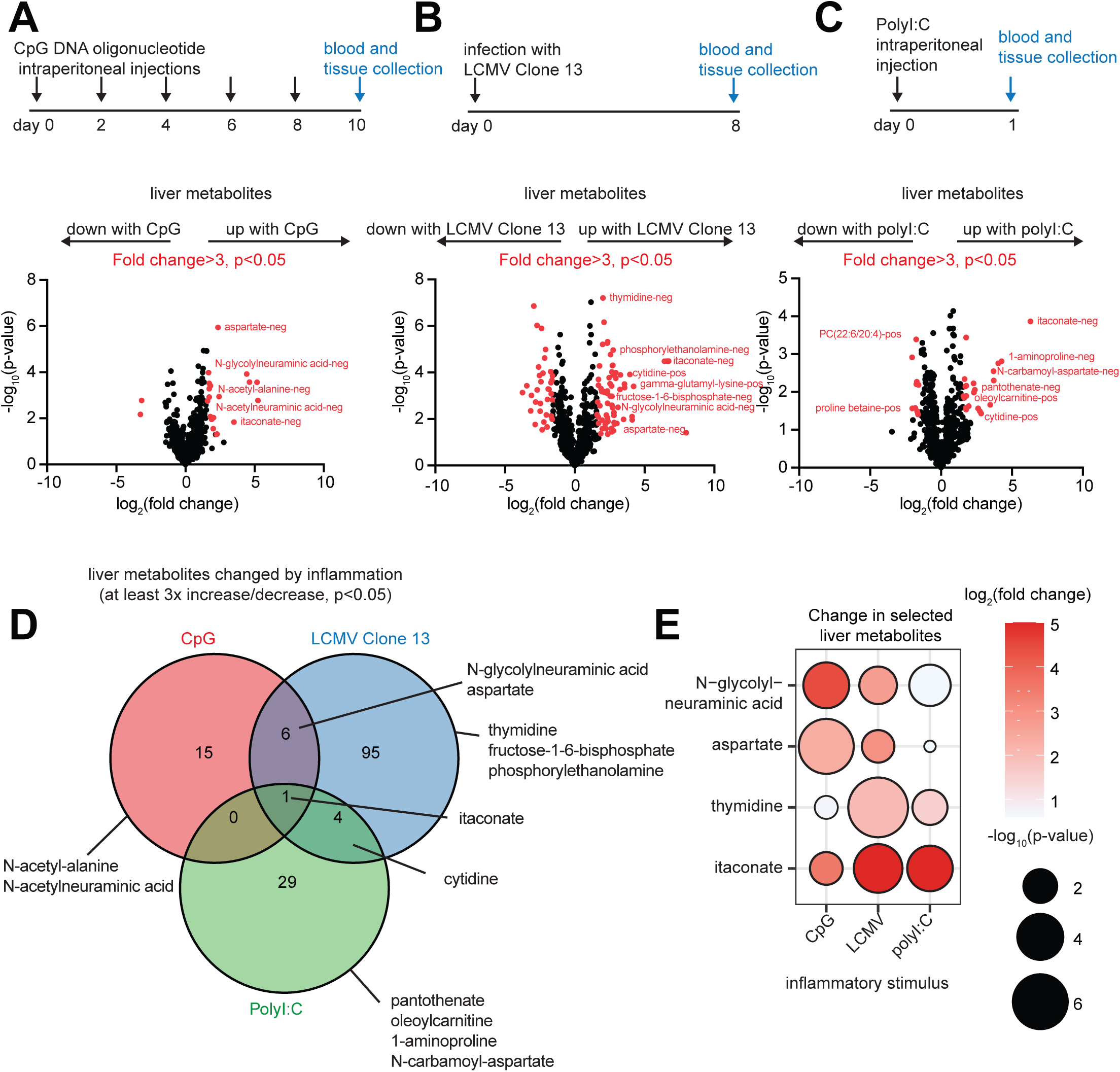
A tissue inflammation metabolomic atlas. A–C: Experimental design and volcano plots of liver metabolites in three mouse models of inflammation: (A) intraperitoneal injection of 50 µg CpG DNA oligonucleotide or control oligonucleotide on days 0,2,4,6,8, with tissue collection on day 10, n=4 mice per group, (B) intravenous infection with 4×10^6^ plaque-forming units of lymphocytic choriomeningitis virus (LCMV) Clone 13, tissue collection on day 8, n=5 mice per group, and (C) 200µg intraperitoneal polyinosinic-polycytidylic acid (PolyI:C) injection or saline injection in controls, tissue collection after 24 hours, n=4 mice per group. Each volcano plot shows log_2_(fold change control versus experimental group) versus -log_10_(p-value) for liver metabolites; metabolites meeting the threshold of greater than 3-fold or less than 0.33-fold change, and p<0.05, are highlighted in red and selected metabolites are labeled. Each panel is representative of two independent experiments, see Supplementary Tables D: Venn diagram of liver metabolites with fold change greater than 3 or less than 0.33, and p<0.05 across the three inflammation models. Selected metabolites are listed. E: Bubble plot showing the log_2_(fold change) and -log_10_(p-value) of selected metabolites across all three inflammatory stimuli. Darker bubble shade is larger log_2_(fold change) in inflammation compared to control, larger size is lower p-value. All p-values were determined using a two-sided t test.

These datasets comprise a tissue atlas of metabolomic changes in inflammation. Across the six tissue types measured (liver, spleen, blood serum, lung, kidney and brain), we found the greatest number of significant changes occurred in the liver for each stimulation (**Figure 1A, Figure S1A-E, Supplementary Tables**). LCMV infection led to the most metabolite changes in the liver (**Figure 1D**), perhaps because ongoing viral replication continually produces pathogen-associated molecular patterns. LCMV altered 107 metabolites in the liver increase or decrease of at least 3-fold and p-value less than 0.05, compared to 22 altered metabolites during CpG-induced cytokine storm and 34 after PolyI:C injection. The majority of changes (66-90%) were metabolite increases in each model, rather than decreases (**Supplementary Tables)**. Surprisingly, the majority of metabolite changes were not shared between the different stimuli (only 10-32% altered liver metabolites shared between stimuli, **Figure 1D**), suggesting that each stimulus elicited a tailored response.

A small set of liver metabolites were changed by two or more stimuli (**Figure 1D-E**). Itaconate was the one metabolite robustly increased across all three *in vivo* immune stimuli. Other shared changes across two out of three conditions included the pyrimidine nucleoside cytidine and the amino acid aspartate, a building block for pyrimidines (**Figure 1A-E, Figure S1F**). Both LCMV infection and CpG cytokine storm dramatically raised the level of a metabolite annotated as N-glycolylneuraminic acid (**Figure 1A-E**, increased 6.5-fold in the liver in LCMV and over twenty-fold in CpG cytokine storm). N-glycolylneuraminic acid is a sialic acid; it is best-studied in its role in protein glycosylation^27^, while the soluble, non-protein-conjugated form has been little-studied.

### Macrophage activation *in vitro* does not replicate most metabolic changes observed in inflamed tissues

Most of the metabolites most altered in inflamed livers, including N-glycolylneuraminic acid, had not previously been studied in their connection with immune response. We hypothesized that perhaps these understudied metabolites were changed in tissues yet not observed in cultured immune cells. To this end, we measured metabolomic changes induced by stimulation of mouse bone-marrow-derived macrophages (BMDM). Macrophages carry a wide variety of sensors for pathogen- and damage-associated molecular patterns^1^, and are a key model system for studying metabolic change in response to immune stimulus. Indeed, itaconate induction and its role in inflammation were first observed in cultured macrophages^16,17^.

We stimulated mouse bone-marrow-derived macrophages with either lipopolysaccharide (LPC), CpG DNA, or PolyI:C (**Figure 2A**), mimicking stimuli present during bacterial and viral infection^28^, and measured metabolomic changes after 24 hours. At this timepoint, we detected robust changes in both metabolite levels and gene expression (**Figures S2A-B**). In contrast to our findings in mouse tissue inflammation, in macrophages different stimuli changed similar metabolites: 73-94% altered metabolites were changed in multiple stimuli in macrophages, versus 32% or less in livers (**Figure 2B-E**). Metabolites increased by two or more stimuli included itaconate; the nucleotide triphosphates ATP and UTP; the glycolytic intermediates fructose-1-6-bisphosphate and dihydroxyacetonephosphate; and the urea cycle intermediate argininosuccinate (**Figure 2E-F**). Enrichment analysis identified many shared altered metabolic pathways across two or all three stimuli, including nucleotide sugars, glycolysis, and the glutathione pathway (**Figure S2C**).

**Figure 2:**
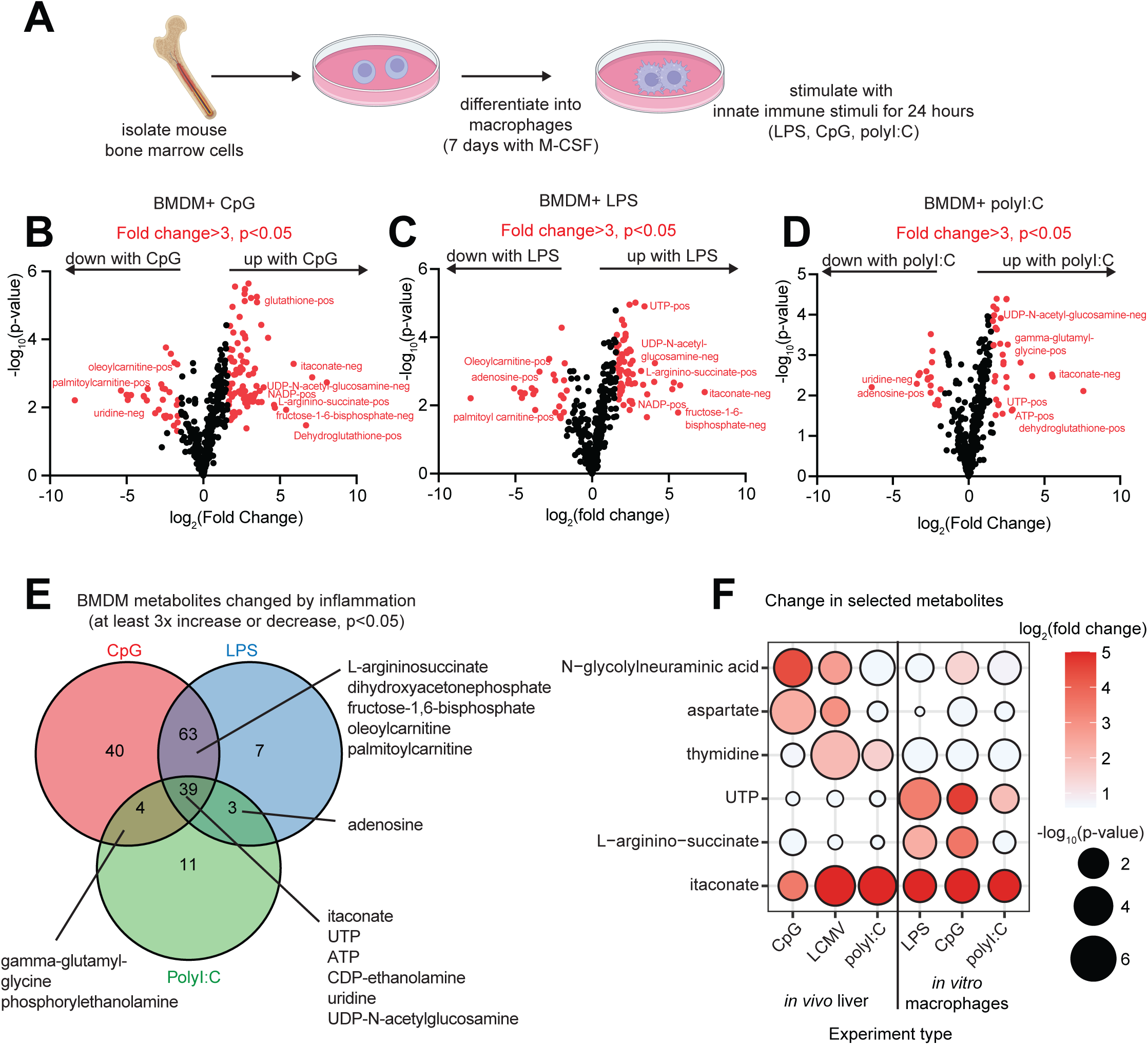
Macrophage activation in vitro does not replicate most metabolic changes observed in inflamed tissues. A: Schematic of *in vitro* bone-marrow-derived macrophage (BMDM) activation. Mouse bone marrow cells were isolated and differentiated into bone marrow-derived macrophages (BMDMs), then stimulated with 4 ng/mL lipopolysaccharide (LPS), 4 ng/mL CpG DNA oligonucleotide, or 500ng/mL polyinosinic-polycytidylic acid (PolyI:C) for 24 hours prior to metabolite extraction. B–D: Volcano plots of BMDM lysate metabolites after 24-hour stimulation with CpG (B), LPS (C), or PolyI:C (D). Metabolites with greater than 3-fold or less than 0.33-fold change and p<0.05 are colored red and selected metabolites labeled. N=3 wells per condition, each experiment is representative of two independent experiments, see Supplementary Figures. E: Venn diagram of BMDM metabolites with greater than 3-fold or less than 0.33-fold change and p<0.05 across LPS, CpG, and PolyI:C stimulations. Selected metabolites shared across conditions are labeled. F: Bubble plot comparing log_2_(fold change) and –log_10_(p-value) of selected metabolites across *in vivo* liver and *in vitro* BMDM inflammatory conditions. Darker bubble shade is larger log_2_(fold change) in inflammation compared to control, larger size is lower p-value. All p-values were determined using two-sided t tests.

With the notable exception of itaconate, most metabolomic changes were not shared between cultured macrophages and mouse liver (**Figure 2F**). For example, UTP was increased 4- to 25-fold in macrophages by different stimuli, but barely changed (0.9- to 1.5-fold) in inflamed livers (**Figure 2F**). Conversely, N-glycolylneuraminic acid, the soluble sialic acid molecule, was induced up to 22-fold in inflamed livers but only up to 2.7-fold in BMDM (**Figure 2F**). These distinct *in vivo* metabolomic changes may explain both why itaconate has been comprehensively studied as a functional immunometabolite^16–19,29^, and why aspartate, thymidine and N-glycolylneuraminic acid have not. These observations also highlight the importance of measuring metabolomic changes in animals and/or human patients.

### Inflammation increases soluble sialic acid in mice and humans

We decided to investigate the regulation and functional role of one of the metabolites induced in tissue inflammation, the soluble sialic acid N-glycolylneuraminic acid (Neu5Gc, **Figure 1**). First, we validated the identity of this compound by confirming that its mass-to-charge ratio, retention time, and tandem-MS fragmentation pattern matched that of a purchased N-glycolylneuraminic acid standard (**Figure 3A**).

**Figure 3:**
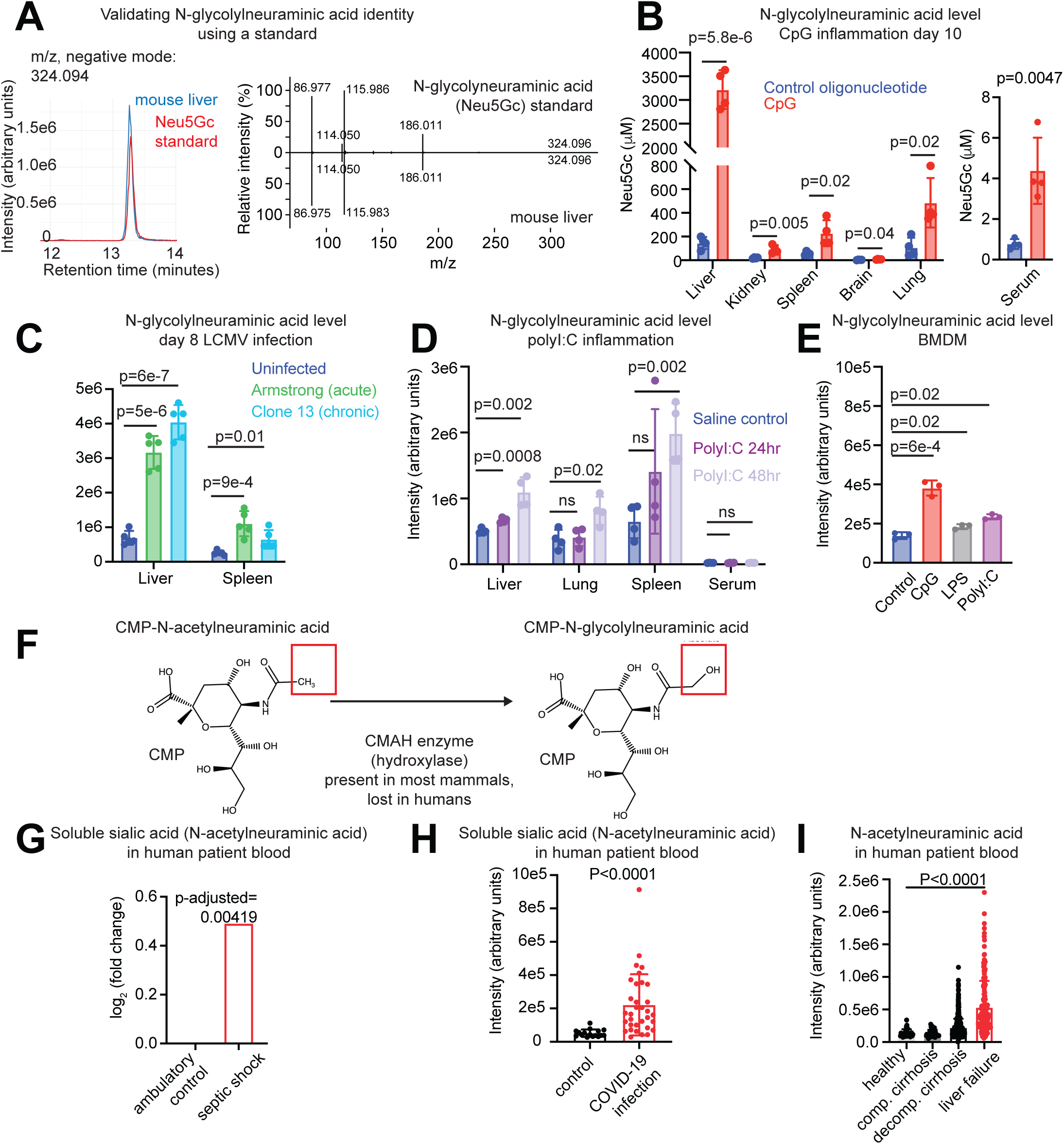
Inflammation increases soluble sialic acid in mice and humans. A: Validation of N-glycolylneuraminic acid (Neu5Gc) identity by comparison of retention time and MS2 fragmentation spectrum of liver peak with a purchased standard (parent m/z 394.0936, negative mode, MS2 collision energy 20V). B: N-glycolylneuraminic acid concentration measured with external standard calibration curve in tissues and blood serum in mice injected intraperitoneally with 50µg CpG DNA or control oligonucleotide on days 0, 2, 4, 6, and 8, with tissue collection on day 10, n = 4 mice per group. C: N-glycolylneuraminic acid level in liver and spleen at day 8 of LCMV infection (either intraperitoneal injection with 2×10^5^ plaque-forming units of Armstrong acute strain or intravenous injection with 4×10^6^ pfu of Clone 13 chronic strain), compared to uninfected controls, n=5 mice per group. D: N-glycolylneuraminic acid level in tissues and blood serum at 24 or 48 hours after 200µg PolyI:C injection, compared to saline-injected controls, n=4 mice per group. E: N-glycolylneuraminic acid level in mouse bone-marrow-derived macrophages after 24 hours of stimulation with 4ng/mL CpG, 4ng/mL LPS, or 500ng/mL PolyI:C, compared to unstimulated controls. Experiments in B-E representative of at least two independent experiments, see Supplementary Figures and Tables. F: Schematic of CMAH hydroxylase reaction to transform CMP-N-acetyl-neuraminic acid to CMP-N-glycolylneuraminic acid; enzyme present in mice but not humans. G–I: Soluble sialic acid (N-acetylneuraminic acid since humans do not produce N-glycolylneuraminic acid) in human patient blood from published inflammation metabolomics datasets. (G) Septic shock versus ambulatory control patients (Rogers et al. 2024^32^). (H) COVID-19 infection versus healthy controls (Thomas et al. 2020^33^). (I) Healthy controls versus patients with compensated cirrhosis, decompensated cirrhosis, or liver failure (Moreau et al. 2020^34^). All p-values from two-sided t tests, except p-adjusted in G calculated in Rogers et al. study.

Next, we asked whether inflammation induced N-glycolylneuraminic acid (Neu5Gc) in tissues beyond the liver. We found that in the CpG cytokine storm model, Neu5Gc was most highly induced in liver, yet was also increased in the blood serum, kidney, spleen, brain and lung (**Figure 3B**). Using an external standard curve we found that Neu5Gc reached extremely high levels in the liver, around 3 millimolar, higher than the level of any amino acid in the blood^30^. LCMV infection also raised N-glycolylneuraminic acid in mouse liver and spleen (**Figure 3C, Figure S3A**). Examining the third model of inflammation we tested, PolyI:C injection, we discovered that even though N-glycolylneuraminic acid was not initially observed as a highly altered metabolite after 24 hours (**Figure 1C-E**), 48 hours after PolyI:C injection N-glycolylneuraminic acid was significantly elevated in the liver and other tissues (**Figure 3D**). An additional inflammatory model, topical application of the TLR7 agonist imiquimod which mimics psoriasis^31^, also induced Neu5Gc in skin, liver and spleen (**Figure S3B**). Therefore, every inflammatory model we have investigated in mice induced the soluble sialic acid Neu5Gc.

In contrast, bone-marrow-derived macrophages stimulated with various TLR ligands only showed a modest induction of this metabolite, between 1.3- and 2.7-fold (**Figure 3E; Figure S3C**). This relatively modest increase of Neu5Gc in macrophages, in contrast to the dramatic induction of metabolites like itaconate and argininosuccinate (up to 137-fold and 12-fold respectively, **Figure 2**), may account for the lack of previous study of Neu5Gc as an immunometabolite.

We next investigated whether soluble sialic acid also increases in human patients experiencing inflammation. N-glycolylneuraminic acid is the most abundant sialic acid in mice and most other mammals, yet this form of sialic acid does not exist in humans, due to a loss of the CMAH enzyme gene^27^ (cytidine monophospho-N-acetylneuraminic acid hydrolase, **Figure 3F**). In humans, the most abundant sialic acid is instead N-acetylneuraminic acid (**Figure 3F**). Publicly available datasets revealed that N-acetylneuraminic acid is elevated in the blood serum in patients with a range of inflammatory conditions, including patients in septic shock^32^ (**Figure 3G**), with COVID-19 infection^33^ (**Figure 3H**), or with liver failure^34^ (**Figure 3I**). Therefore, soluble sialic acid is an evolutionarily-conserved metabolite elevated in inflammation.

### Soluble sialic acid is produced from sialylated proteins by CD11b+ myeloid cells in the liver

Since we found that soluble sialic acid is increased in a variety of inflammatory states in both mice and humans, we asked how this molecule is produced. There has been extensive study of the synthesis of protein-bound sialic acids^27,35^, but the soluble form is much less studied. Protein-bound sialic acids are produced from a pathway including the intermediates UDP-N-acetyl-glucosamine and CMP-sialic acid (**Figure 4A**), then sialyltransferase enzymes add the sialic acid moiety from CMP-sialic acid onto a protein-conjugated glycan chain as the terminal sugar^27^. Soluble Neu5Gc is produced if neuraminidase enzymes (Neu1-4) cleave sialic acid off a glycosylated protein^36^. However, the vast production of Neu5Gc we observed in inflammation, over 3mM in the liver during CpG cytokine storm (**Figure 3B**), required that we confirm that this conventional pathway was used and not some previously-unknown route.

**Figure 4:**
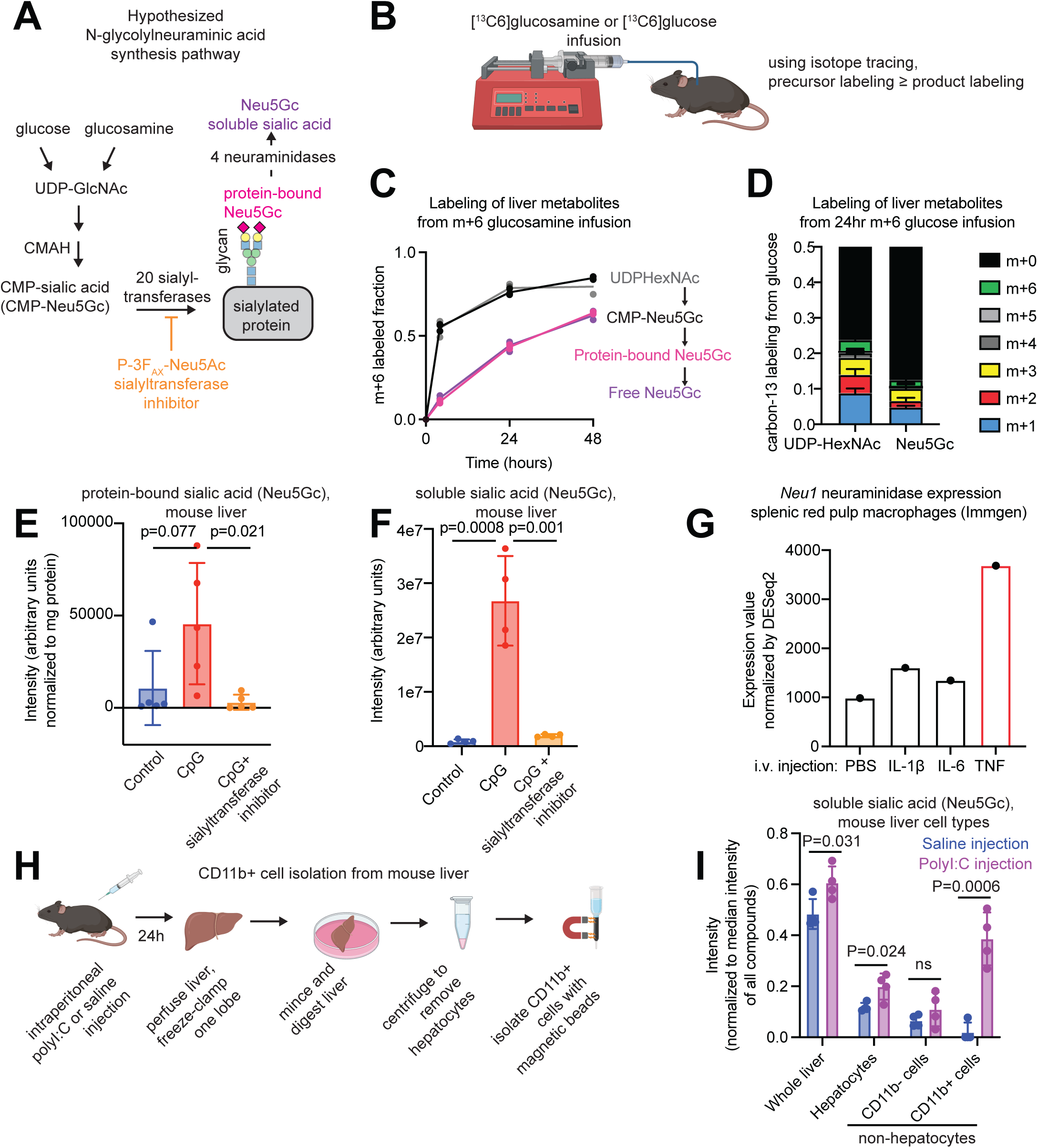
Soluble sialic acid (N-glycolylneuraminic acid) is produced from sialylated proteins by CD11b+ myeloid immune cells in the liver. A: Hypothesized synthesis pathway for soluble N-glycolylneuraminic acid. Glucose or glucosamine is used to produce UDP-GlcNAc, which is converted to CMP-Neu5Gc and incorporated into glycoproteins by a sialyltransferase enzyme. A neuraminidase subsequently cleaves glycans to release soluble N-glycolylneuraminic acid. B: Schematic of isotope tracing by continuous intravenous infusion of carbon-13-labeled substrates into awake, catheterized mice. Labeling enrichment of any precursor metabolite is always equal or greater than that of its product metabolites. C: Mass+6 (m+6) fractional carbon-13 labeling of UDP-HexNAc, CMP-Neu5Gc, protein-bound N-glycolylneuraminic acid measured by hydrolysis from isolated protein, and free soluble Neu5Gc in mouse liver after continuous infusion of m+6 ^13^C-glucosamine (m+6) for 4, 24 or 48 hours. N=3 mice for 4 and 24 hours, n=2 mice for 48 hours. D: Carbon-13 isotopologue distribution (m+0 through m+6) for UDP-HexNAc and soluble Neu5Gc after 24 hours of m+6 ^13^C-glucose infusion, n=3 mice. E: Protein-bound N-glycolylneuraminic acid measured by protein isolation from mouse livers and hydrolysis with acetic acid, in mice injected with control oligonucleotide, with CpG, or with both CpG and 30mg/kg P-3F_AX_-Neu5Ac sialyltransferase inhibitor over 10 days, n=4 mice per group. F: Same liver samples from E, soluble N-glycolylneuraminic acid measured by LC-MS metabolomics. G: Expression of *Neu1* neuraminidase RNA in splenic red pulp macrophages in mice intravenously injected with the listed cytokine or PBS, data from Lee et al. 2025^39^. H: Schematic of cell type isolation from mouse liver using digestion and CD11b+ magnetic bead selection. I: Soluble N-glycolylneuraminic acid level normalized to the median intensity of all metabolites in whole liver, isolated hepatocytes, CD11b− non-hepatocytes, and CD11b+ cells from mouse livers 24 hours after 200µg polyI:C or saline intraperitoneal injection, n = 4 mice per group. All p-values from two-sided t-tests. All experiments are representative of 2 independent experiments, see Supplementary Figures.

To confirm that the described pathway is used to produce Neu5Gc in mouse tissues, we first used heavy-isotope-labeled nutrient intravenous infusion (schematic **Figure 4B**). The upstream precursor of sialic acids, UDP-N-acetyl-glucosamine (UDP-GlcNAc), can be produced from glucose or from glucosamine^37^ (**Figure 4A**). Therefore, if we infuse carbon-13-glucose or - glucosamine, any metabolite made from UDP-GlcNAc would therefore have similar or less carbon-13 labeling as UDP-GlcNAc (**Figure 4B**). Consistent with the previously-described pathway, Neu5Gc in the liver of healthy mice indeed displayed carbon-13 labeling from intravenously-infused carbon-13-glucosamine as well as glucose (**Figure 4C-D, Figure S4A-B**). The labeling was of a reduced magnitude compared to the precursors UDP-HexNAc and CMP-Neu5Gc, which is likely due to the large pool size of protein-bound sialic acid. These tracing experiments are consistent with soluble Neu5Gc being produced from protein-bound Neu5Gc as expected (**Figure 4A**).

We next wondered how N-glycolylneuraminic acid level is increased during inflammation. Two non-mutually-exclusive possibilities are that more sialylated protein is produced during inflammation, or that there is increased neuraminidase activity during inflammation. To address these possibilities, we measured the level of protein-conjugated as well as free N-glycolylneuraminic acid in livers of mice after 10-day induction of CpG cytokine storm (**Figure 4E-F**). Although there is a trend toward increased protein-bound sialic acid in livers of CpG-injected mice, with the mean value increased two- to four-fold (**Figure 4E, Figure S4C**), the increase in free Neu5Gc in CpG-treated mice is around twenty-fold (p=0.0008, **Figure 4F**). Therefore, increased neuraminidase activity is the major contributor to the increase in liver Neu5Gc in inflammation.

To further confirm that Neu5Gc in inflammation was produced by cleavage from sialylated protein, we treated mice with the sialyltransferase inhibitor P-3F_AX_-Neu5Ac along with CpG cytokine storm induction. P-3F_AX_-Neu5Ac is expected to prevent sialic acid addition to glycosylated proteins^38^, and our data confirmed that this inhibitor indeed lowered liver protein-bound sialic acid (**Figure 4E**). If soluble Neu5Gc were produced by cleavage from sialylated proteins, reducing the sialylated protein pool with sialyltransferase inhibitor should reduce soluble Neu5Gc. This was indeed the case: treating mice with the sialyltransferase inhibitor during CpG cytokine storm reducing liver soluble Neu5Gc close to the level in uninflamed mice (**Figure 4F**). Therefore, soluble N-glycolylneuraminic acid is produced via sialyltransferase conjugation to proteins followed by neuraminidase cleavage.

Since the liver contains a mix of cell types including immune cells, fibroblasts, and hepatocytes, we asked what is the cellular source of Neu5Gc in the liver. We observed in publicly available data^39^ that Neuraminidase-1 (*Neu1*) RNA is induced in splenic macrophages *in vivo* by TNF (**Figure 4G**), and not in other immune cells. Since TNF is induced by CpG and polyI:C (**Figure S4B**)^24^, its stimulation of *Neu1* RNA levels suggests that myeloid cells might be responsible for the increased cleavage and release of Neu5Gc during inflammation. Consistent with this hypothesis, macrophages upregulate neuraminidase activity during differentiation^40^ and activation^41^.

To definitively test whether macrophages produce Neu5Gc during inflammation *in vivo*, we performed metabolomics on isolated hepatocytes, as well as CD11b+ and CD11b- non-hepatocyte cells purified with magnetic beads from livers of mice injected with PolyI:C (**Figure 4H-I, Figure S4D-G**). Consistent with our previous data, N-glycolylneuraminic acid was barely induced in whole liver 24 hours after PolyI:C injection (**Figure 1E**, **3D**, **4I**). Strikingly, we found that N-glycolylneuraminic acid was robustly increased in liver CD11b+ myeloid cells (**Figure 4I, Figure S4D-G**), though it was undetectable in liver myeloid cells of saline-injected controls. Therefore, we hypothesize that during inflammation, cytokines including TNF induce neuraminidase expression in myeloid cells, leading these cells to produce high levels of soluble Neu5Gc by sialylated protein cleavage.

It is unclear why soluble Neu5Gc is only modestly induced when macrophages are stimulated in tissue culture (**Figure 3E, Figure S3A**), compared to the dramatic induction in myeloid cells *in vivo* (**Figure 4I**). One possibility is that macrophages rely on another cellular source such as hepatocytes to produce abundant glycosylated protein, which macrophages take up and cleave, while in culture sialylated protein is less abundant.

### N-glycolylneuraminic acid does not affect pathology in cytokine storm or viral infection

The metabolite itaconate is highly induced in myeloid cells during inflammation or infection, and exerts important antimicrobial and immune-regulatory roles^18,19,29^. We asked whether Neu5Gc, also highly induced in myeloid cells during inflammation, also plays a functional role. To address this question, we took advantage of the sialyltransferase inhibitor P-3F_AX_-Neu5Ac, which blocked the induction of soluble Neu5Gc during CpG-induced inflammation (**Figure 3E-F, Figure S5A**). In fact, we found that this drug reduced Neu5Gc with moderate specificity, only reducing three other metabolites more than 3-fold (**Figure 5A**).

**Figure 5:**
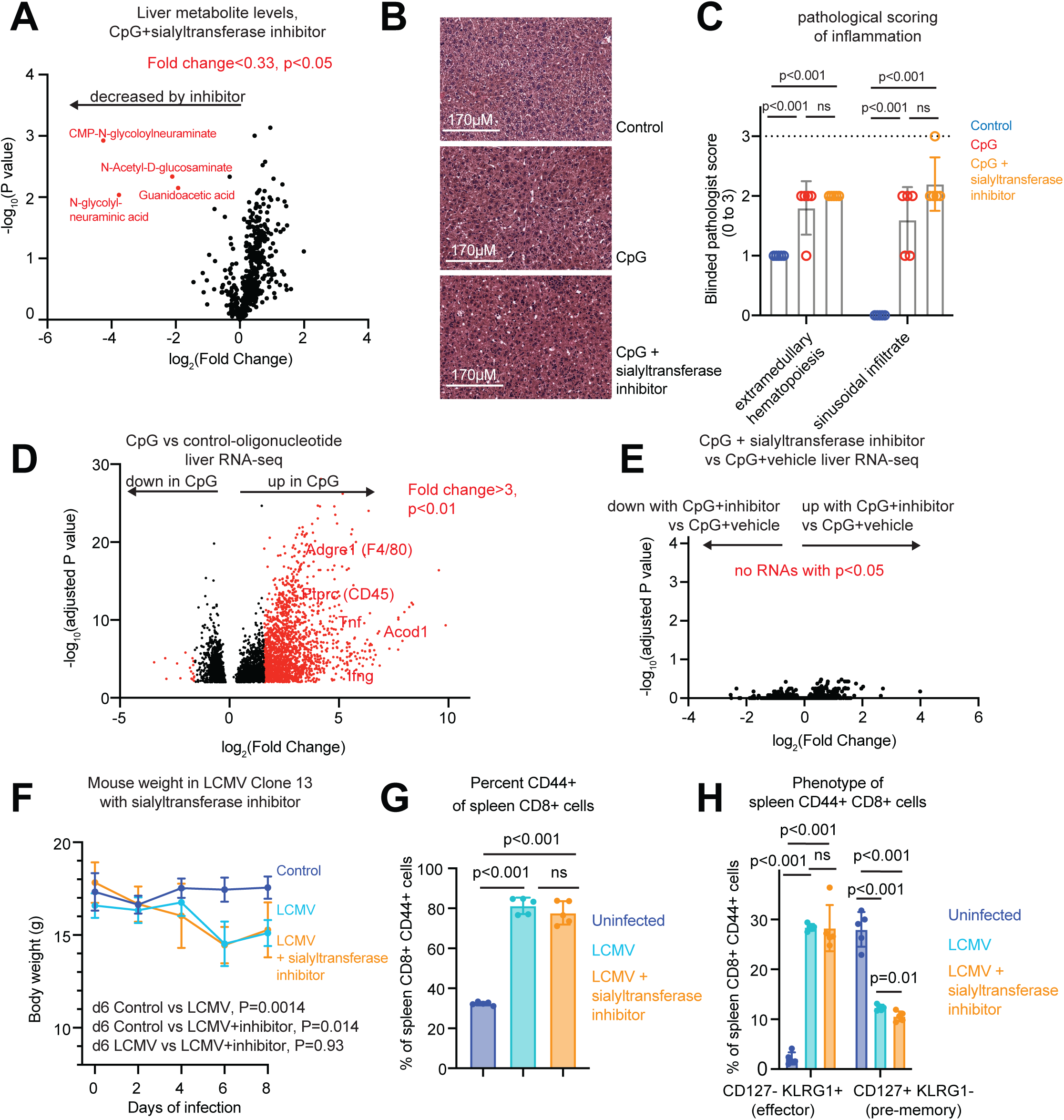
N-glycolylneuraminic acid does not affect pathology in cytokine storm or viral infection. A: Volcano plot of liver metabolites from mice after 10 days of every-other-day CpG injection, with daily 30mg/kg P-3F_AX_-Neu5Ac sialyltransferase inhibitor or vehicle control co-treatment, n=4 mice per group. Metabolites in red have fold change less than 0.33 and p<0.05. B: Representative H&E-stained liver sections from control oligonucleotide injected, CpG plus vehicle, and CpG plus sialyltransferase inhibitor-treated mice. C: Blinded liver histopathology scoring in control, CpG, and CpG plus sialyltransferase inhibitor-treated mouse livers. Zero is feature not detected, 3 is most extreme manifestation of feature. D: Volcano plot of liver RNA-sequencing comparing CpG- to control-oligonucleotide-injected mice. RNAs with fold change less than 0.33 or greater than 3 and Benjamini-Hochberg adjusted p-value less than 0.01 are plotted in red, and selected RNAs are labeled. Large number of RNAs with adjusted p>0.01 are not plotted. E: Volcano plot of liver RNA-sequencing comparing mice injected with CpG and vehicle, or CpG and P-3F_AX_-Neu5Ac sialyltransferase inhibitor. No RNAs were changed with Benjamini-Hochberg adjusted p<0.05. F: Mouse body weight over 8 days in LCMV-Clone-13-infected mice, infected mice treated daily with P-3F_AX_-Neu5ac, and uninfected controls, n=5 mice per group. P values compare weights from day 6 using two-sided t-tests. G: Percentage of CD44+ cells of splenic CD8+ T cells in uninfected controls, LCMV-Clone-13-infected mice, and LCMV-Clone-13-infected mice treated with sialyltransferase inhibitor, measured by flow cytometry, n=5 mice per group. H: Percentage of CD44+ CD8+ splenic T cells exhibiting effector (CD127− KLRG1+) and pre-memory (CD127+ KLRG1−) phenotypes, measured by flow cytometry, n=5 mice per group. All p values from two-sided t-tests, except Benjamini-Hochberg adjusted p-values in 5D-E determined by DESeq2.

However, reducing Neu5Gc levels did not seem to alter liver pathology in CpG-induced cytokine storm. Using blinded histopathological assessment, increased foci of extramedullary hematopoiesis, increased prominence of cells lining the sinusoids, and increased cellular infiltrates within the sinusoids were identified in livers with CpG administration compared to control animals (**Figure 5B-C, Figure S5B**). However, daily treatment with sialyltransferase inhibitor neither worsened nor improved these phenotypes. While CpG administration induced a number of RNAs in the liver including *Tnf*, *Ifng,* and *Acod1* (the enzyme that produces itaconate) (**Figure 5D**, **Figure S5C**), there were *no changes* in RNA level comparing livers of mice receiving CpG alone versus CpG with sialyltransferase inhibitor (**Figure 5E**). Therefore, blocking Neu5Gc induction did not alter gene expression or pathology during CpG cytokine storm.

We next investigated whether blocking Neu5Gc induction might alter pathology due to LCMV Clone 13 chronic viral infection, and found no evidence that it did. The Clone 13 strain of LCMV causes mice to lose weight due to the antiviral immune CD8+ T cell response^3^, but blocking Neu5Gc production with sialyltransferase inhibitor did not ameliorate or exacerbate this weight loss (**Figure 5F, Figure S6A).** We used flow cytometry to interrogate the CD8+ T cell response on day 8 of viral infection and found that sialyltransferase inhibitor had no detectable effect (**Figure 5G-H, Figure S6B**). For example, the fraction of spleen CD8+ T cells that were activated (CD44^hi^) was increased by LCMV infection as expected, yet was not changed by sialyltransferase inhibitor (**Figure 5G**). Similarly, the proportions of activated CD8+ T cells displaying effector (KLRG1+ CD127-) or precursor-like (CD127+KLRG1-) phenotypes were unchanged or minimally changed by the drug (**Figure 5H**).

Further, we asked whether high levels of Neu5Gc could directly interfere with viral infection, since sialic acid is an entry receptor for many viruses including Influenza A virus^42^. We conducted an Influenza A infection assay in the Calu-3 lung epithelial cell line, in the presence or absence of soluble sialic acids (either Neu5Gc or the dominant human form Neu5Ac, **Figure S6C**). Addition of sialic acid did not alter rates of Influenza A virus infection. Therefore, so far we have found no functional role for soluble sialic acid in infection or inflammation, although it is potently increased in both human and mouse inflammation.

## Discussion

Here, we provide an atlas of metabolomic changes in inflammation in mouse models and in tissue culture macrophages. We identified metabolites increased during immune responses *in vivo*, including N-glycolylneuraminic acid, aspartate, and thymidine. We then characterized the production route, cellular source, and function of one of these novel immunometabolites, the soluble sialic acid N-glycolylneuraminic acid (Neu5Gc). We showed that this metabolite is induced in mice by all *in vivo* inflammatory stimuli tested. The analogous human metabolite, N-acetylneuraminic acid, was also induced by inflammation including sepsis, COVID-19 infection, or liver failure. We showed that Neu5Gc is produced by cleavage and release of protein-bound sialic acid by CD11b+ myeloid cells. However, we could find no effect of Neu5Gc production on pathology, gene expression, viral infection, or T cell phenotype. Our experiments demonstrate that metabolomic changes in mice or human patients during immune activation are distinct from changes induced in tissue culture macrophage activation.

It was surprising to us that most metabolite changes in inflamed tissues were distinct from changes in activated macrophages. A notable exception was itaconate, which increased in inflamed tissues and macrophages (**Figure 1**–**2**); as expected, itaconate was produced *in vivo* by myeloid cells (**Figure S4G**). Why were other metabolite changes not conserved between cultured macrophages and inflamed tissues? First, cell types other than myeloid cells may be dominant producers of specific metabolites; for example our previous work suggested that thymidine may be produced by T cells during inflammation^45^. Note however that even in healthy tissues, myeloid cells are by far the most abundant immune cell type^46^, and their numbers rise during inflammation; therefore these cells may well be dominant contributors to metabolic changes in inflamed tissues in many cases. Second, there may be signals or metabolic substrates present *in vivo* that are not replicated in tissue culture. This second possibility may be the explanation for the stronger induction of Neu5Gc *in vivo*, since myeloid cells produce it in the inflamed liver (**Figure 4H-I**) but not in culture when activated with the same stimulus (**Figure 3E**).

Identifying which cell type is responsible for production of any metabolite *in vivo* is technically challenging, since tissues in mice and humans are a mix of different cell types. Here, our data supports a role for CD11b+ myeloid cells in the production of Neu5Gc soluble sialic acid (**Figure 4, Figure S4**). (Note that several cell types, including macrophages and neutrophils, express the CD11b+ marker^47^; so far we don’t know which of these cell subsets may be the dominant Neu5Gc producer.) However, a caveat of our cell isolation experiments is that tissue digestion and cell purification can alter metabolite levels, since the cells live and metabolize in an altered nutrient milieu during the isolation procedure^15,48,49^. The definitive way to test this hypothesis would be to generate a mouse conditionally deficient for Neu5Gc production in myeloid cells- for example LysM-cre Neu1^fl/fl^ ^50^- and test whether Neu5Gc is no longer induced when the mouse is exposed to an inflammatory stimulus. Another approach would be to use imaging mass spectrometry to colocalize tissue Neu5Gc to myeloid cells^51^, although currently the spatial resolution of this technique precludes single-cell resolution.

Why would this metabolite Neu5Gc be so highly induced- up to 3mM in CpG cytokine storm liver-yet play no functional role? Possibly Neu5Gc does play a role in a context not examined here: for example, in the resolution of inflammation or wound healing. Alternatively, it is possible that neuraminidase activity itself, rather than the metabolite it produces, can play a role in inflammation, perhaps via its role in removing sialic acid from glycosylated proteins. Sialylated glycoproteins send immunosuppressive signals via the family of Siglec receptors^52^; neuraminidase removal of the terminal sialic acid from glycoproteins could help unleash the immune response. Our experiments using P-3F_AX_-Neu5Ac blocked the addition of sialic acid onto glycoproteins; yet perhaps Neu1 cleavage off mature glycoproteins has a distinct effect on the immune response.

Overall, we demonstrate that tissue immune responses elicit a distinct set of changes compared to macrophage activation in culture, and therefore the cellular sources and biological consequences of metabolic change in inflamed tissues deserves deeper study.

## Methods

### Mouse strains

Mouse experiments were approved by the University of Pennsylvania Institutional Animal Care and Use Committee. Experiments used C57Bl/6 mice aged 9-12 weeks obtained from Charles River Laboratory. C57BL/6 mice infected with LCMV were 7 weeks old and obtained from Jackson Laboratory.

### CpG injection model of inflammation

CpG oligodeoxynucleotides were obtained from Invivogen (ODN 1826, item number tlrl-1826) and reconstituted in sterile saline (0.25mg/ml). Control oligodeoxynucleotide was also obtained from Invivogen (ODN 2138, item number tlrl-1826c-1). Mice received intraperitoneal injections of 200µl of CpG ODN or control ODN (50 µg) on days 0, 2, 4, 6 and 8, with tissues generally collected on day 10, unless otherwise specified.

### PolyI:C model of immune response

Mice received intraperitoneal injection of 200ug of PolyI:C (polyinosinic-polycytidylic acid, Invivogen, item number tlrl-pic) reconstituted in 100 µl of sterile saline, while control mice were injected with 200 µl saline. Blood and tissues were collected 24 or 48 hours after injection.

### Lymphocytic choriomeningitis virus infection

LCMV Armstrong and Clone 13 were grown in BHK cells (American Type Culture Collection (ATCC), CL-10) and titrated by performing plaque assays on Vero cells (ATCC, CCL-81) as described previously^53^. Mice were infected intraperitoneally with LCMV Armstrong (2 × 10^5^ plaque-forming units (p.f.u.)) or intravenously via the tail vein with LCMV Clone 13 (4 × 10^6^ p.f.u.) diluted in 1% FBS/RPMI. Mouse body weight was measured daily, and blood and tissues were collected for metabolomics and flow cytometry after 8 days.

### Imiquimod psoriasis model

Mouse backs were shaved, then 60mg 5% imiquimod cream (Perrigo) or 60mg Vaseline as control was applied daily for 7 days. Tissues were collected on day 7.

### P-3F_AX_-Neu5Ac drug treatment

P-3F_AX_-Neu5Ac sialyltransferase inhibitor^38,54^ was obtained from BioTechne (item number 5760), and reconstituted in 10% DMSO and sterile saline. Mice received 30mg/kg of P-3F_AX_-Neu5Ac daily via intraperitoneal injection of 200µl for the duration of the study, either 8 days for LCMV infection or 10 days for CpG stimulation. Control groups received injection 200µl of 10% DMSO in saline.

### Jugular vein catheterization

Using aseptic surgical techniques, a catheter (INSTECH item number C10PU-MCA2A10) was inserted in the right jugular vein of the mouse. The catheter was connected to a vascular access button (INSTECH item number VABM1B/25) implanted under the skin on the back of the mouse. After surgery, all mice recovered for at least 3 days before infusion of tracers.

### [U-^13^C_6_]glucosamine infusion to measure labeling of metabolites

Jugular-vein-catheterized mice (described above in jugular vein catheterization) received intravenous infusion of 50mM [U-^13^C_6]_ glucosamine (Omicron Biochemicals Inc., item number GLC-091) reconstituted in sterile saline at 0.1microliter/minute/ gram body weight for 4, 24 or 48 hours via a programmable syringe pump (New Era Syringe Pumps, item number: NE-1000). Mice were infused in their own cages and were able to move freely while attached to infusion line. Water and standard chow were available ad libitum, and 7am to 7pm light-dark cycle was maintained during the infusion as in the housing room.

### [U-^13^C_6_] glucose infusion to measure labeling of metabolites

Nutrient infusion was carried out in awake freely-moving mice for 24 hours, as described above for [U-^13^C_6_] glucosamine, with the following changes. Mice were infused with 800mM of [U-^13^C_6_] glucose (Cambridge Isotope Laboratory item number CLM-1396-PK) reconstituted in sterile saline. The infusion started at 6pm at 0.0375 µl/min/g for 1 hour. From 7pm to 7am the rate was 0.1ul/min/g, it then returned to 0.0375 µl/min/g from 7am to 6pm. Water and standard chow were available ad libitum, and 7am to 7pm light-dark cycle was maintained during the infusion as in the housing room.

### Serum sampling for measurement of metabolite levels or carbon-13 enrichment

Blood was collected by snipping the tail tip of the mouse and squeezing downward into a Sarstedt microvette blood tube (Thermo Fisher, item number NC9059691) and immediately placed on ice. Blood samples were then centrifuged at 4°C for 10 minutes at 21,300RCF and serum was transferred to a 1.5ml microcentrifuge tube (item 3499, Thermo Fisher Scientific). Serum was stored at -80°C until processed.

### Tissue sampling for carbon-13 metabolite enrichment measurements after nutrient infusion

Mice were euthanized by cervical dislocation and tissues were quickly collected and immediately freeze-clamped using a Wollenberger clamp cooled in liquid nitrogen. This approach minimizes metabolic changes, in particular due to hypoxia^55,56^. All tissues were stored at -80°C until processed.

### Tissue sampling for metabolomics measurements (with no nutrient infusion)

Prior to harvesting tissues, uninfused mice were fasted for 6 hours (approximately 9am-3pm) by transferring to a fresh cage with no food, to standardize the feeding state across mice. The mice were euthanized by cervical dislocation and tissues were collected, placed in labeled pieces of aluminum foil, and immediately freeze-clamped using a Wollenberger clamp cooled in liquid nitrogen. All tissues were stored at -80°C until processed.

### Tissue sampling for flow cytometry after LCMV infection

The mice were euthanized by cervical dislocation and tissues were clamped as above. Part of the spleen was freeze clamped, and lymphocytes were isolated from the remaining unclamped spleen by mincing and pushing through a 70-μm nylon filter (Thermo Fisher, item number 08-771-2). Resulting single-cell suspensions were treated with ACK Lysing Buffer (Thermo Fisher item number A1049201) for 3 min at room temperature before washing cells with 5% FBS/RPMI.

### Separation of liver cells for metabolomics and flow cytometry after PolyI:C injection

For liver cell separation and analysis by flow cytometry 24 hours after PolyI:C injection, the livers of saline or polyI:C-injected mice were flushed in situ prior to dissection, by injecting sterile saline via the inferior vena cava and drained from the liver by cutting the hepatic portal vein. 1ml of saline was flushed through the liver before a lobe was collected and placed in MACS buffer (PBS, Thermo Fisher, item number: MT21040CM, 2mM Trypsin-EDTA, Thermo Fisher, 25200114, 0.5% BSA, Sigma Aldrich, 3117057001). At the same time a lobe was freeze-clamped for whole-liver metabolomic measurement.

The non-clamped liver lobe was finely minced with scissors and digested in 5mL of 1 mg/mL collagenase (Thermo Fisher, item number NC1839860) and 50 U/mL DNAse I (Millipore Sigma, item number 10104159001) in MACS buffer for 30 min at 37C degrees while shaking at 180 RPM. The digest was passed through a 100uM filter (Thermo Fisher, item number 08-771-19).

The cell suspension was spun at 20 RCF for 5 minutes at 4°C to pellet hepatocytes. The supernatant was collected and further centrifuged at 450 RCF for 5 min at 4°C. The pellet from the second spin was resuspended in 900 µl MACS buffer, to which 100 µl of anti-Cd11b (Miltenyi Biotec, item number 130-126-725) beads were added. This was incubated for 10 minutes at 4°C. After incubation, a further 4 mL of MACS buffer was added. Cell suspensions were then applied to magnetic columns (Miltenyi item 130-042-401), placed in magnet (Miltenyi item 130-091-051), with flow-through analyzed as CD11b- cells. Then column was removed from magnet and MACS buffer pushed through to elute CD11b+ cells. All fractions of cells were collected for flow cytometry and mass spectrometry analyses.

### Mouse bone-marrow-derived macrophage differentiation and stimulation

1×10^7^ bone marrow derived macrophages were thawed and resuspended in 20ml of macrophage growth media (RPMI, Thermo Fisher, item number: SH30027FS, + 20% heat inactivated FBS, Sigma Aldrich, item number: F2442500ML, 20ng/ml recombinant M-CSF, Thermo Fisher, item number: 315-02-50UG, 1% pen/strep, Emsco Fisher, item number: SV30010) and divided between two 10cm non-tissue culture treated plates (Emsco Fisher, item number: 08-757-100D). 10ml of fresh growth media was added on the fourth day of culture.

After 7 days in growth media, macrophages were detached from plates by washing with PBS+2 millimolar EDTA, and then counted. Cells were centrifuged 10min at 300 RCF 4°C, resuspended in RPMI+10% heat-inactivated FBS+10ng/mL M-CSF at a ratio of 1 million cells per milliliter media. Cells were plated at 1e6/ml in 1.5ml media per well in 6-well plates. Cells were allowed to settle for 6-12 hours, and then immune stimuli were added. For CpG ODN 1826 (Invivogen item number tlrl-1826, stock solution 1µg/ml) and LPS (Sigma item number L3024, stock solution 1µg/ml), 6 µl was added per well for a final concentration of 4ng/mL. For PolyIC (Invivogen, item number tlrl-pic, stock solution 50µg/mL), 15µl was added to each well for a final concentration of 0.5µg/mL. All treatments were applied for 24 hours (or other length of time where indicated) before cells were collected for LC-MS analysis.

### Influenza infection experiments in cultured cells

Human adenocarcinoma lung epithelial cells, Calu-3 (ATCC HTB-55), were cultured in MEM (GIBCO 11095-072) supplemented with 10% (v/v) fetal bovine serum (FBS) (Corning 35-015-CV), 1% (v/v) penicillin/streptomycin (Gibco 15140-122), 1% (v/v) Glutamax (Gibco 35050-061) and 1% (v/v) non-essential amino acid (Gibco item number 11140-050), in a 37°C, 5% CO_2_ incubator. 4×10^5^ cells were plated in 12 well collagen coated tissue culture plate (Corning 356500). Cells were pre-treated with vehicle, N-acetylneuraminic acid (VWR, item number 80055-610), N-glycolylneuraminic acid (Sigma Aldrich, item number G9793-25MG) and Baloxavir (MedChemExpress S-0033447) at the concentration indicated for two hours. Following two-hour pre-treatment cells were infected with Influenza virus A (PR8 strain; courtesy Dr. Scott Hensley) for 24 hours at the indicated multiplicity of infection.

### Blood serum and tissue processing and extraction for metabolite measurements

Serum was extracted for MS analysis by the addition of ice-cold 100% methanol, 50 times the volume of serum on ice. The samples were vortexed, then left to precipitate overnight at - 80°C. They were then centrifuged at 4°C, for 25 minutes, at 21,300RCF. The supernatant was transferred to a MS vial.

For tissue metabolite extractions, tissues were kept frozen on dry ice or in pre-cooled metal racks on dry ice at all times. Tissue samples were placed in pre-cooled locking 2ml tubes (Emsco Fisher, item number: 05-402-95) with a metal ball (Retsch item 224550003) and powdered using the CryoMill (Retsch item number 207490001). Powdered tissues were then quickly weighed into a new pre-cooled 1.5ml tube and weights were recorded, with target amount of powder between 5-20mg per sample. A volume of methanol: acetonitrile: water in a 2:2:1 ratio, acidified with 0.5% formic acid^57^ (v/v), was added to the tube 40 times the weight of the powder in microliters (acetonitrile, VWR BDH83639.400; water VWR BDH23595.400, methanol Fisher Scientific A4544). Tubes were vortexed and then 8% v/v ammonium bicarbonate (Sigma Aldrich, item number 09830-500G, 15%weight/volume in water) was added, before vortexing again. Samples were left to precipitate on dry ice for 10 minutes before centrifuging at 4°C, 25 minutes, 21,300 RCF. Supernatants were transferred to a new tube, and the centrifugation step repeated. Then, samples were transferred to MS vials (vials, Thermo Scientific, 6ERV11-03PPC; caps, Thermo Scientific, 6PRC11ST1R) for LC-MS measurement.

Cells from liver digestion and cell isolation were extracted in 100µl of methanol:acetonitrile:water (2:2:1 by volume) and kept on dry ice. After vortexing, they were centrifuged at 4°C, for 25 minutes, at 21,300RCF. The supernatant was then transferred to a MS vial for analysis.

### Macrophage extraction for metabolite measurements

For metabolite extraction from bone-marrow-derived macrophages cultured in six-well-plates, media was aspirated, then 300 microliters of dry-ice-cooled 80% methanol in water was quickly added to each well. Plate was swirled to spread solvent across the whole well, then plate was placed on metal surface over dry ice for 15 minutes. After 15 minutes, solvent from each well was transferred to corresponding 1.5ml plastic tube and stored in -80°C freezer. After storage overnight or longer to enable protein precipitation, samples were centrifuged at 4°C, for 25 minutes, at 21,300RCF. Supernatant was transferred to a second tube and again centrifuged at 4°C, for 25 minutes, at 21,300RCF. Supernatant from the second spin was transferred to an MS vial for analysis.

### Measurement of protein-bound Neu5Gc

To measure protein-bound sialic acid, liver samples were collected and powdered as described above. Powders were weighed and weights were recorded. To all powdered samples 400µl of methanol, 200µl of chloroform (Emsco Fisher, item number: AA32614K2) and 300µl of water were added before centrifuging. The upper aqueous layer containing free soluble metabolites was discarded. The protein layer (between the aqueous and organic layers) and the lower organic layer was retained. Methanol was added to the remaining sample, the sample was centrifuged and the supernatant discarded 2 more times before air drying for 15 minutes. To the remaining protein pellet, 100µl of glacial acetic acid was added. Samples were heated to 80 degrees, shaking, for 6 hours before being dried down under nitrogen manifold. 100µl of methanol was added to the tubes, and centrifuged. The supernatant was transferred to MS vials for analysis. When analyzed, peak intensity was normalized to mass of the input tissue powder.

### Histology scoring

Lobes of liver were collected and immediately washed in PBS. They were placed in individual cassettes (Fisher Scientific, item number B851739BL) and transferred to ice cold 4% paraformaldehyde (Electron Microscopy Sciences item 15710). Samples were kept at 4°C rocking for 72 hours. After 72 hours, cassettes were rinsed in PBS and stored in 50% ethanol.

The veterinary pathologists performing the histopathological analysis are part of the University of Pennsylvania Penn Vet Comparative Pathology Core Facility (RRID:SCR_022438). Hematoxylin and eosin-stained slides of formalin fixed, paraffin embedded liver tissue were evaluated by an ACVP board-certified veterinary pathologist. The tissues were evaluated using a semiquantitative method. Parameters scored included necrosis (0 – no evidence of necrosis, 1 – rare single cell necrosis, 2 – small clusters of necrotic cells, 3 – multifocal necrotic foci which extend greater than one hepatic lobule), extramedullary hematopoiesis (EMH) (0 – no evidence of EMH, 1 – scattered small clusters of EMH, less than 1 per hepatic lobule on average, 2 – multifocal EMH foci, between 1 and 3 per hepatic lobule on average, 3 - multifocal EMH, greater than 3 foci per hepatic lobule on average) and prominence of sinusoidal lining cells with sinusoidal infiltrate (0 – no evidence of increased cellularity within sinusoids, 1 - mild sinusoidal cell prominence, 2 - moderate sinusoidal cell prominence and increased cellularity in sinusoids, 3 - marked sinusoidal cell prominence and increased cellularity in sinusoids).

### LC-MS metabolomics measurement

Metabolites in samples prepared as described above were measured on the Q-Exactive Plus Orbitrap mass spectrometer (Thermo Fisher) which was coupled to a Vanquish Horizon Liquid Chromatography system (Thermo Fisher). A Waters XBridge BEH Amide Column (item number 186006724) with Waters BEH Amide XP VanGuard Cartridge (item number 186007763) was used. Mobile phase A was water:acetonitrile 95:5 (v/v) with 20mM ammonium acetate (Sigma Aldrich, item number: 09689) + 20mM ammonium hydroxide (Sigma Aldrich, item number: 09859), pH 9.45, with 5uM (NH_4_)_2_HPO_4_ (Thermo Fisher, item number: AC201822500) and 50mM medronic acid (Millipore Sigma, 1984-15-2). Mobile phase B was acetonitrile. Flow rate was 0.15ml/min throughout and the gradient over the course of 25 minutes went as follows: 0 min, 90%B; 2 mins, 90%B; 3 minutes, 75% B; 7 minutes, 75% B; 8 minutes, 70% B; 9 minutes, 70% B; 10 minutes, 50% B; 12 minutes, 50% B; 13 minutes, 25% B; 14 minutes, 25% B; 16 minutes, 0% B; 20.5 minutes, 0% B; 21 minutes, 90% B; 25 minutes, 90% B.

Injections were 5 µl in volume, and both negative and positive scans were performed over a mass range of 70 to 1000 m/z, with a resolution of 70,000. Automatic gain control was set at 3e6. Source ionization parameters were optimized with the spray voltage at 4.0 kV and in-source CID of 5.0eV, and other parameters were as follows: capillary temp at 425, S-lens level at 50, aux gas heater temperature at 400, sheath gas at 35, aux gas at 10, and sweep gas flow at 2. Data was collected by the Thermo Fisher software Xcalibur (v4.5.474.0).

### Measurement of soluble Neu5Gc concentration in blood serum and tissues

A 1mM Neu5Gc (Cayman Chemicals, item number 30283) stock solution was made in water, then diluted to concentrations of 500, 400, 300, 200, 100 and 10μM to make a calibration curve. Calibrators were then extracted in 2:2:1 methanol:acetonitrile:water and run alongside mouse tissue and serum samples. Concentration was calculated by fitting a line to the calibrators and calculating the unknown concentrations in tissues and serum samples.

### MS/MS analysis

A 1mM Neu5Gc (Cayman Chemical, item number 30283) stock solution was made in water, then extracted with 2:2:1 methanol: acetonitrile: water, with a 1:50 ratio of stock solution:solvent. MS/MS analysis was performed on the Q Exactive Plus using the liquid chromatography conditions described above. Mass spectrometry parameters were as described earlier but scanning only negative mode and with resolution of 140,000. Additionally, parallel reaction monitoring was performed for analytes with an m/z of 324.09360 (with a 2m/z window). Collision energy was set to 20V, AGC to 2e6 and resolution was 17,500.

### LC-MS data analysis

LC-MS data analysis was performed using El-Maven (v0.12.1) using an in-house knowns list with metabolite masses and retention times; identities were assigned with a tolerance of +/-15.00ppm and +/- 1 minute in retention time; peak areatop was used to determine intensities. When analyzing enrichment from carbon-13-labeled nutrient infusion, the natural abundance of carbon-13 was corrected for using the Accucor (v0.3.0) package in R^58^.

### Flow cytometry

For LCMV spleens after ACK lysis (see Tissue sampling for flow cytometry after LCMV infection above), extracellular antibody staining was performed using the following antibodies: anti-CD44-BUV395 BD Biosciences item 740215, anti-CD8b-BUV563 BD Biosciences item 741342, anti-B220-BUV615 BD Biosciences item 569724, Zombie Yellow fixable viability kit Biolegend item 423103, anti-NKG2A/C/E-BUV661 BD Biosciences item 741582, anti-CD127-BUV737 BD Biosciences item 564399, anti-KLRG1-BUV805 BD Biosciences item 741993, anti-CD62L-BV605 Biolegend item 104437, anti-CX3CR1-BV650 Biolegend item 149027, anti-CD39-BV711 BD Biosciences item 567295, anti-Vα2-BV750 BD Biosciences item 747258, anti-Tim3-BV785 Biolegend item 119725, anti-CD94-FITC Biolegend item 105506, anti-Ly108-BB700 BD Biosciences item 742272, anti-Lag3-PE/Dazzle594 Biolegend item 369217, anti-CD69-Cy5 Biolegend item 104509, anti-PD1-PeCy7 Biolegend item 109109. Then cells were fixed and permeabilized using the Foxp3 kit (Thermo Fisher item 501128857). Intracellular staining was then performed using the following antibodies: anti-Tox APC Miltenyi item 130-107-785, anti-granzyme B BV421 BD Bioscience item 563389, anti-Tcf1-PE Cell Signaling Technologies item 14456, and anti-Ki67-AF700 BD Biosciences item 561277. Analysis was performed on the BD LSRFortessa X50. Data was analysed in FlowJo v10.

After isolation of CD11b+ and CD11b- cells from livers of PolyI:C treated mice (see Separation of liver cells for metabolomics and flow cytometry after PolyI:C injection above), cells were aliquoted into 96 well plates and centrifuged at 1800 rpm for 2 mins at 4°C. Live Dead Violet (Biolegend, 423113) master mix was diluted 1:500 in phosphate buffered saline and added to all wells. A positive control and compensation well was prepared by heat-killing cells at 65c for 5 mins and combining in a 1:1 ratio with live cells before staining with Zombie violet (Biolegend, item number 423113). The plate was incubated at room temperature in the dark for 15 minutes before spinning, washing with MACS buffer and adding antibodies. Antibodies were as follows: CD45 BV605 – 610/20 YG, CD11b FITC -515/20 -Blue, F4/80 APC - 670/30 Red, Zombie violet - 423 Violet (Biolegend, item numbers 103155, 123116, 101206, 423113). All antibodies were diluted 1:300. The plate incubated at 4°C for 20 minutes, then was washed twice in MACS buffer. Analysis was performed on the BD FACSymphony. Data was analysed in FlowJo v10.

### Gene expression analysis

RNA-sequencing was performed by Azenta Genewiz. RNA for sequencing was isolated from powdered liver tissues using Quick-RNA MicroPrep Kit from Zymo Research (item number R1051).

Analysis of liver samples or macrophage samples by qPCR was achieved by extracting RNA with Quick-RNA MicroPrep Kit from Zymo Research (item number R1051). cDNA was produced using the iScript™ cDNA Synthesis Kit (Bio-Rad, item number: 1708891). Primers for qPCR analysis were all purchased from IDT and are outlined in the table. qPCR was performed using the QuantStudio 6 Pro (Thermo Fisher) with PowerUp SYBR green (Thermo Fisher, item number: 4309155). Normalization to *Tbp* RNA was used across all samples and primers.

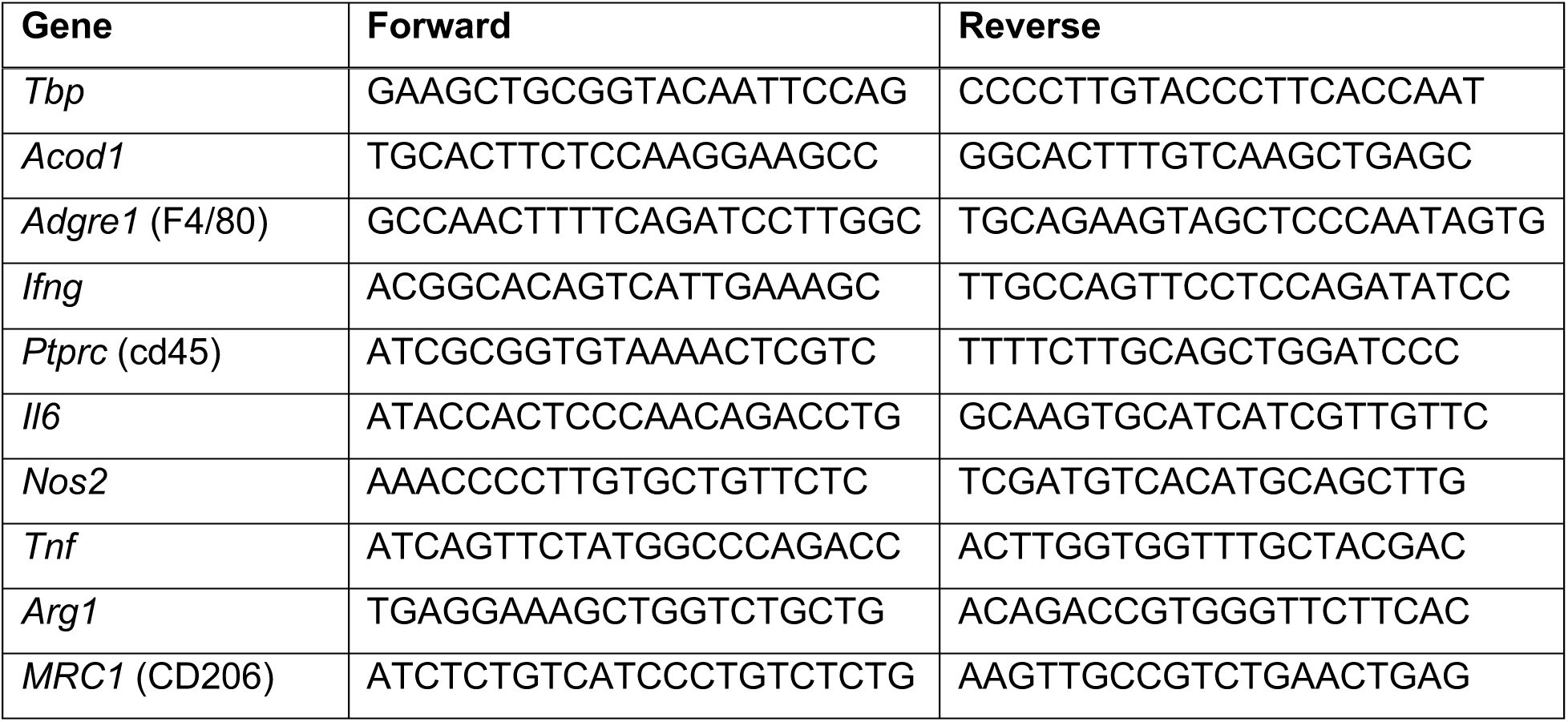

For measurement of Influenza A virus RNA, Calu-3 cells infected by Influenza A virus (see above) were lysed in TRIzol, and RNA was extracted using on-column purification (Zymo Research #R1018) with DNase I treatment (Zymo Research #E1010). cDNA was generated from random hexamers primers to prime reverse transcription reactions with Moloney murine leukemia virus (M-MLV) reverse transcriptase (Ambion). The qPCR was performed in with the diluted cDNA using a SYBR-Green based (ThermoScientific) qPCR assay on AppliedBiosystems equipment. Relative quantification was calculated according to the standard curve method by normalization to the ribosomal 18s loading control. Fold change was calculated by normalizing to the vehicle control.

### Statistical analysis

All p values calculated by two-sided t-tests. Fold changes and t-test p-values for volcano plots were calculated in R.

### Metabolic pathway enrichment analysis

All significantly altered compounds (log_2_fold-change>1.58, p-value<0.05) from a given experiment were entered into “Enrichment Analysis” (“Over Representation Analysis” tab) in MetaboAnalyst^59^. The Small Molecule Pathway Database metabolite library was selected for the reference metabolome, and we only considered pathways with at least 2 metabolite members. Then significantly enriched pathways were manually checked for those where at least 3 metabolites in the pathway were significantly changed in the dataset, and the enrichment made logical sense (e.g. in some cases the same altered metabolites like ATP were considered members of many different pathways in which case not all results were listed).

## Figure construction

All figures were constructed in GraphPad Prism, except for Venn diagrams and bubble plots made using ggplot2^60^ in R, schematics were created using BioRender.

## Acknowledgements

This work was supported by R00CA273517 and V Foundation V2024-009 to CRB. Thanks to Mikel Haggadone, Mark Boyer, and Sunny Shin for help with the bone-marrow-derived macrophage culture system, and thanks to Josh Rabinowitz for feedback in starting the project.

## Supplementary Figures

**Figure S1, related to Figure 1:**
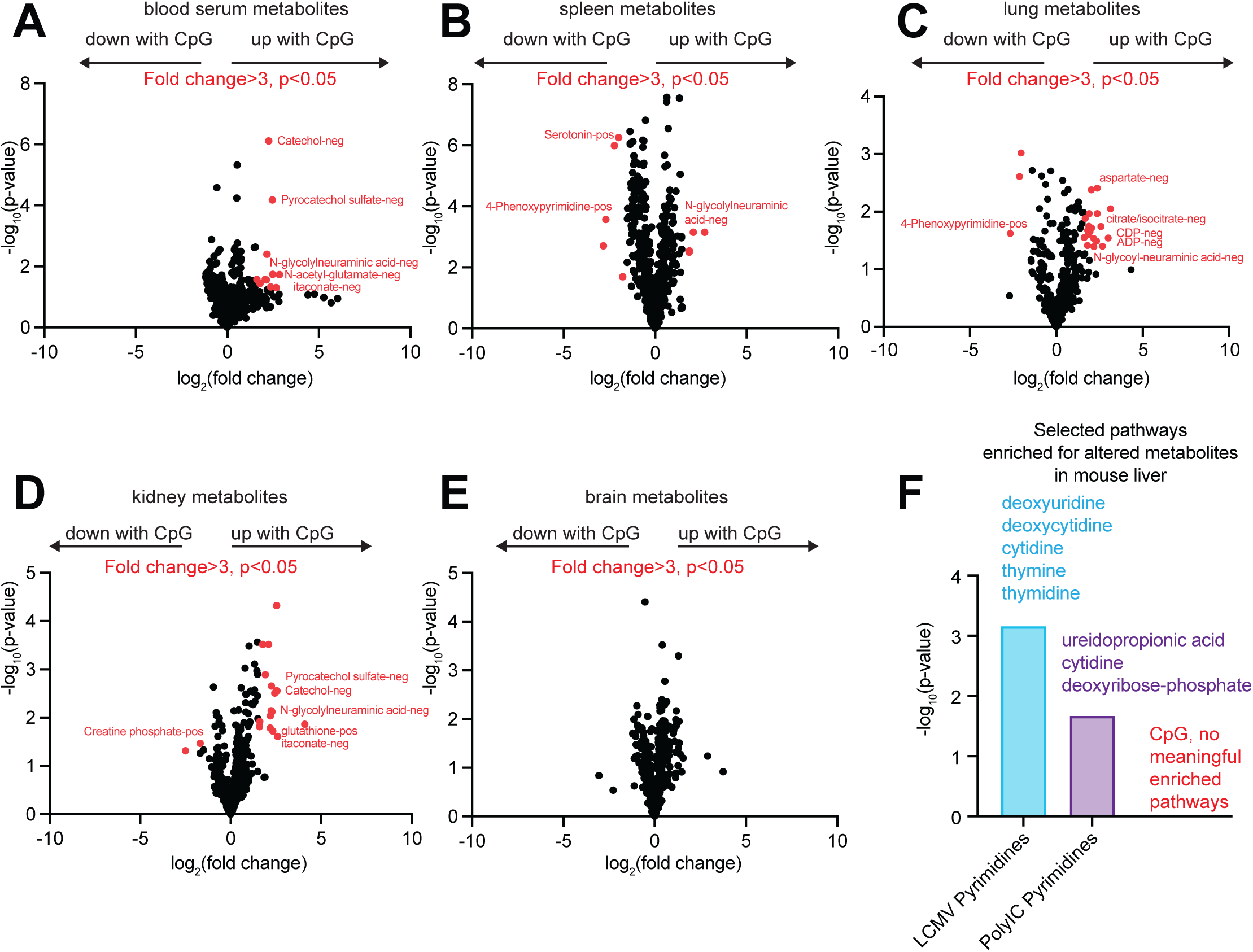
Metabolomic changes and pathway enrichments in inflamed mouse tissues. A-E: Metabolites changed between control mice and mice receiving intraperitoneal injection of 50 µg CpG DNA oligonucleotide (or control oligonucleotide) on days 0,2,4,6,8, with tissue collection on day 10, n=4 mice per group. (A) metabolites in blood serum, (B) spleen, (C) lung, (D) kidney, (E) brain; metabolites meeting the threshold of greater than 3-fold or less than 0.33-fold change, and p<0.05, are highlighted in red and selected metabolites are labeled, p-values were determined using a two-sided t test. (F) Pathways enriched for altered metabolites in mouse liver after 8 days of LCMV Clone 13 infection, 24 hours after polyI:C injection, or after 10 days of CpG injection; data corresponds to Figure 1. All enriched pathways with more than 2 metabolites represented are shown.

**Figure S2, related to Figure 2:**
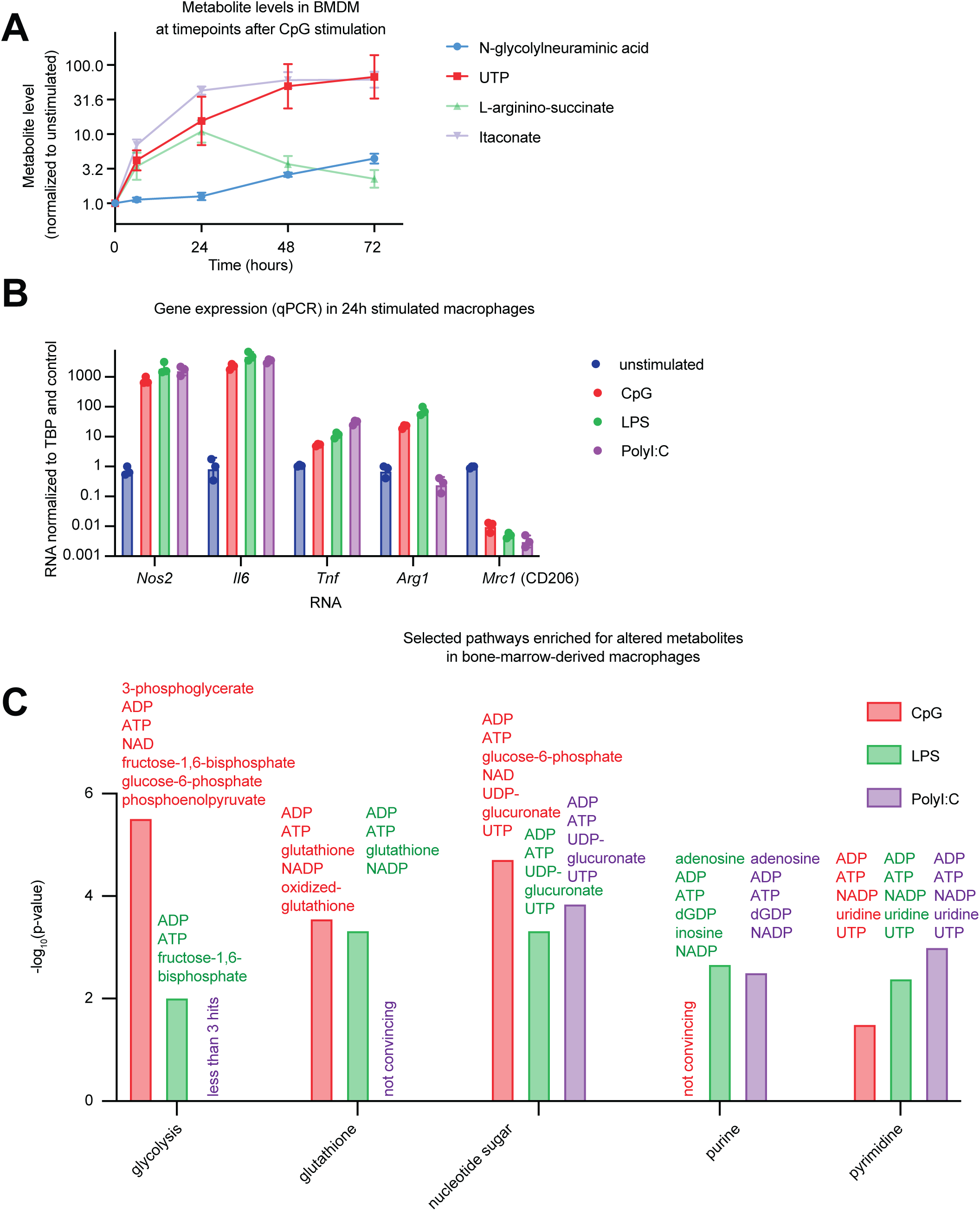
Metabolomic changes, gene expression changes, and pathway enrichments in stimulated bone-marrow-derived macrophages. A: Changes in selected metabolites at 0,4,24,48 or 72 hours after stimulation with 4ng/mL CpG oligonucleotide, showing that 24 hours is an early timepoint where large metabolite changes are detected. B: Quantitative PCR for selected genes in bone-marrow-derived macrophages 24 hours after stimulation with 4 ng/mL CpG DNA oligonucleotide, 4 ng/mL lipopolysaccharide (LPS), or 500ng/mL polyinosinic-polycytidylic acid (PolyI:C). C: Pathways enriched for altered metabolites in macrophages after 24 hours of stimulation with CpG, LPS or PolyI:C, data corresponds to Figure 2. Selected enriched pathways with more than 2 metabolites represented.

**Figure S3, related to Figure 3:**
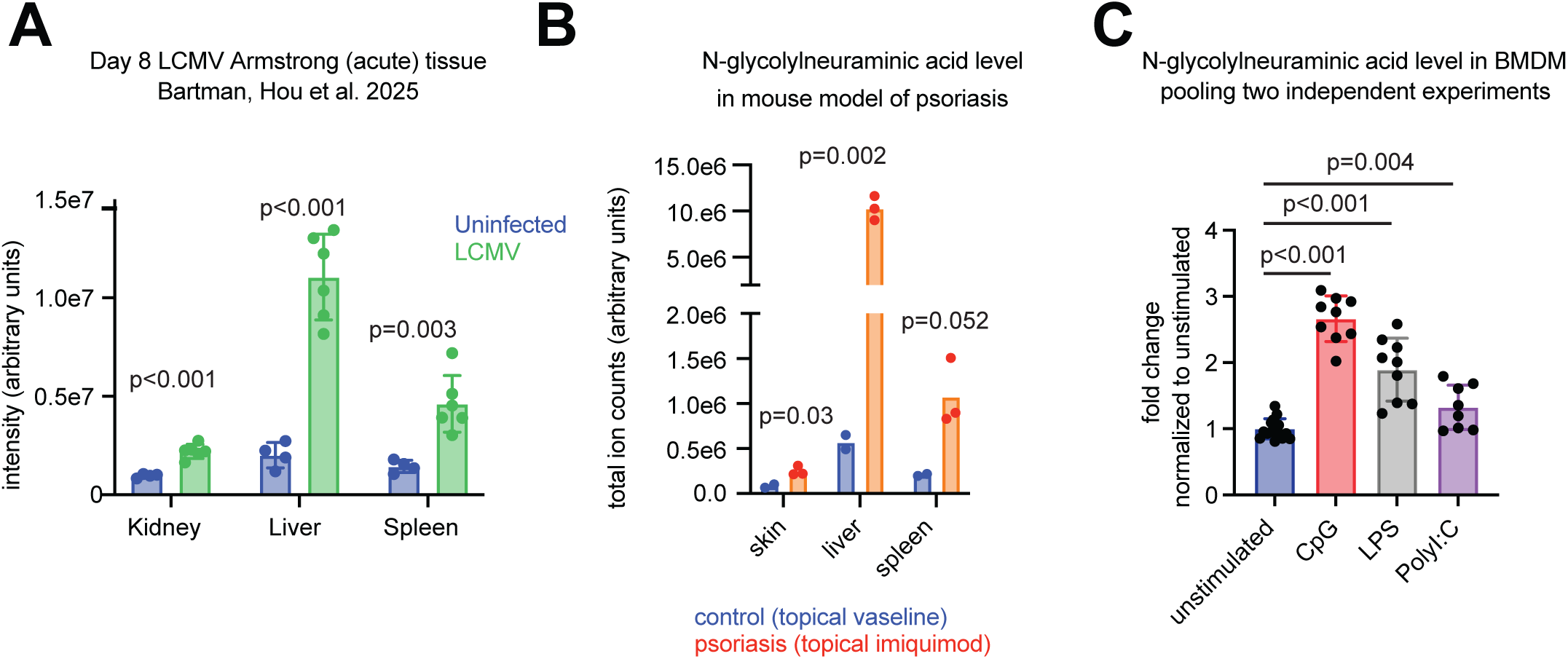
Inflammation increases N-glycolylneuraminic acid more in mouse tissues than in cultured macrophages. A: N-glycolylneuraminic acid level from kidney, liver and lung of mice infected with acute Armstrong LCMV for 8 days, n=4 uninfected and n=6 infected mice, data from Bartman, Hou et al. 2025^45^. B: N-glycolylneuraminic acid level in mouse skin, liver and spleen induced by 7 days topical application of 60mg imiquimod or vaseline control, n=2 control and n=3 imiquimod-treated mice. C: N-glycolylneuraminic acid level in bone-marrow-derived macrophages 24 hours after stimulation with 4 ng/mL CpG DNA oligonucleotide, 4 ng/mL lipopolysaccharide (LPS), or 500ng/mL polyinosinic-polycytidylic acid (PolyI:C), normalized to unstimulated macrophages from same experiment. All p-values from two-sided t-tests.

**Figure S4, related to Figure 4:**
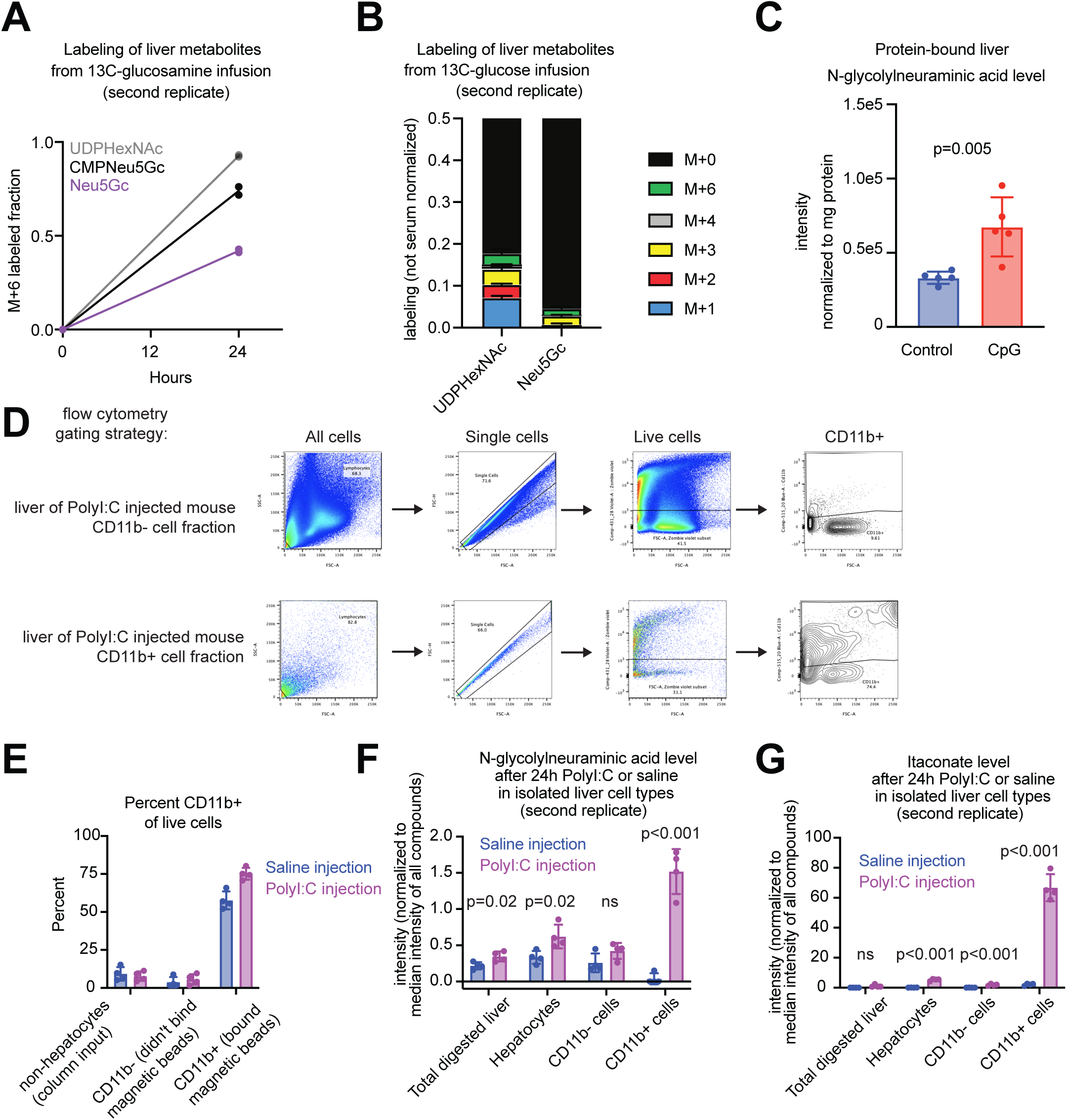
N-glycolylneuraminic acid is produced from sialylated proteins by CD11b+ myeloid immune cells *in vivo*. A : Mass+6 (m+6) fractional carbon-13 labeling of UDP-HexNAc, CMP-Neu5Gc, and free soluble Neu5Gc in mouse liver after continuous infusion of m+6 ^13^C-glucosamine (m+6) 0 or 24 hours, n=1 uninfused and n=2 24h-infused mice. B: Carbon-13 isotopologue distribution (m+0 through m+6) for UDP-HexNAc and soluble Neu5Gc after 24 hours of m+6 ^13^C-glucose infusion, n=2 mice. C: Protein-bound Neu5Gc measured by protein isolation from mouse livers and hydrolysis with acetic acid, in mice injected with control oligonucleotide or with CpG over 10 days, or with both CpG and 30mg/kg P-3F_AX_-Neu5Ac sialyltransferase inhibitor, n=5 mice per group. D-E: Flow cytometry data to assess purity of liver CD11b+ myeloid cell isolation, related to data in Figure 4I. F: N-glycolylneuraminic acid level normalized to the median intensity of all metabolites in whole liver, isolated hepatocytes, CD11b− non-hepatocytes, and CD11b+ cells from control and PolyI:C-treated mice, n = 4 mice per group, second independent replicate of experiment in Figure 4I. G: Itaconate level normalized to the median intensity of all metabolites in whole liver, isolated hepatocytes, CD11b− non-hepatocytes, and CD11b+ cells from control and PolyI:C-treated mice, n = 4 mice per group. As expected, itaconate is enriched in CD11b+ myeloid cells. All p-values from two-sided t-tests.

**Figure S5, related to Figure 5:**
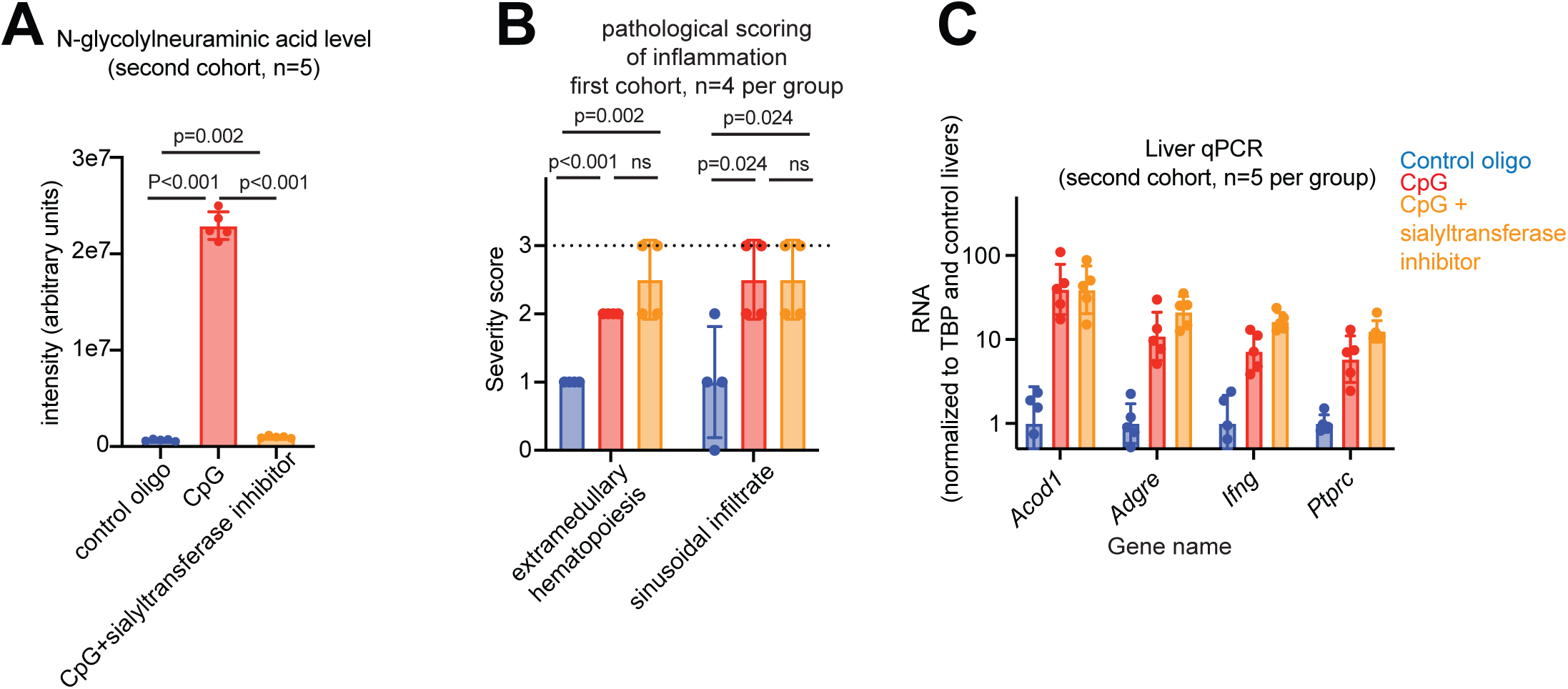
N-glycolylneuraminic acid does not affect pathology in CpG-induced cytokine storm. A : Soluble N-glycolylneuraminic acid level in livers of mice injected over 10 days with control oligonucleotide, CpG oligonucleotide, or CpG plus daily 30mg/kg P-3FAX-Neu5Ac sialyltransferase inhibitor injection, n=5 mice per group. B: Blinded pathologist scoring of liver inflammation in mice injected over 10 days with control oligonucleotide, CpG oligonucleotide, or CpG plus 30mg/kg P-3FAX-Neu5Ac sialyltransferase inhibitor, n=4 mice per group, second independent replicate of experiment in Figure 5C. C: Quantitative PCR of selected RNAs in livers of mice from Figure S5A, n=5 mice per group. in All p-values from two-sided t-tests.

**Figure S6, related to Figure 5:**
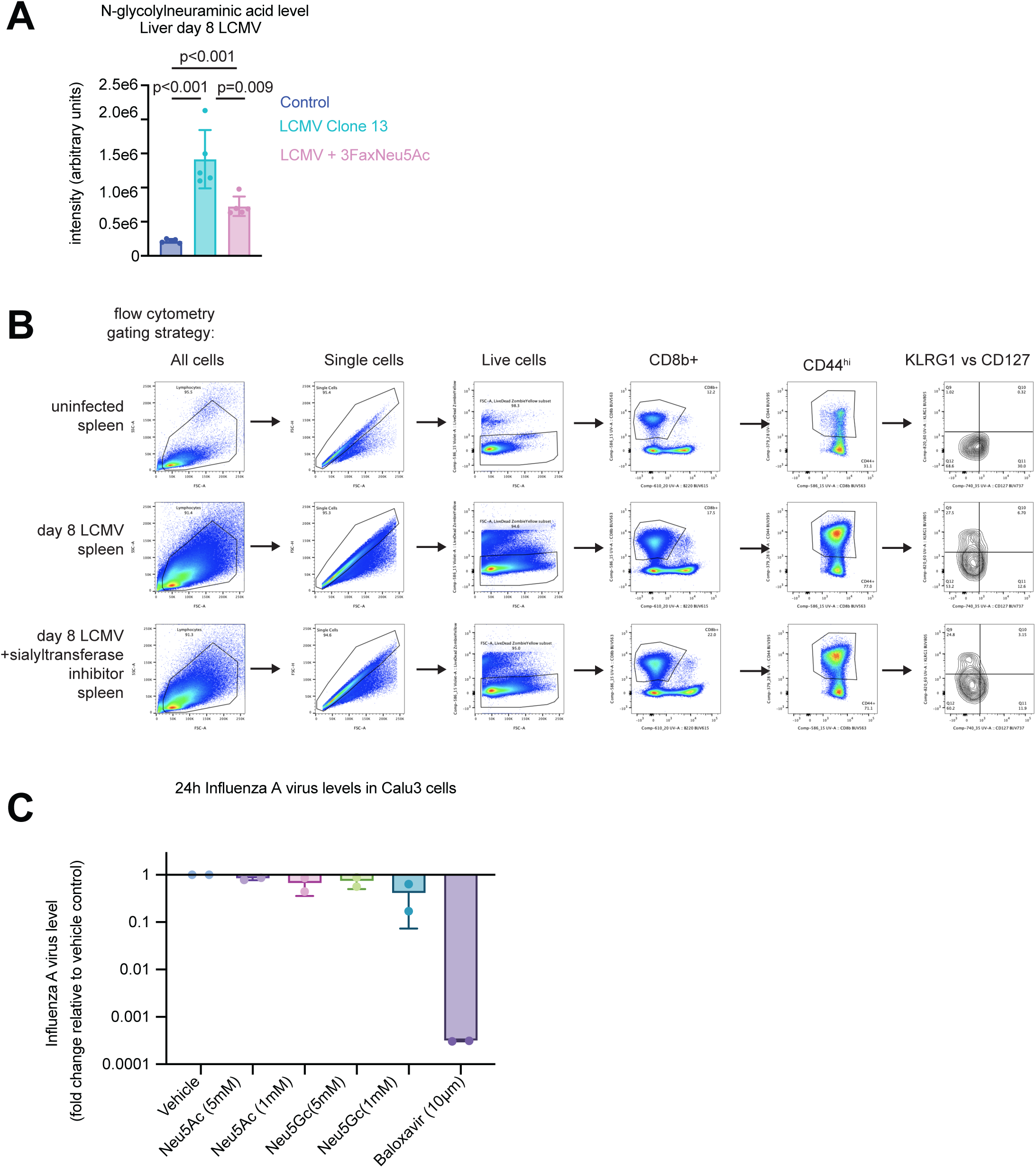
N-glycolylneuraminic acid does not affect viral infection. A: Soluble N-glycolylneuraminic acid level in livers of uninfected mice, mice 8 days after LCMV Clone 13 infection, or LCMV clone 13 plus daily 30mg/kg P-3FAX-Neu5Ac sialyltransferase inhibitor injection, n=5 mice per group, p-values from two-sided t-tests. B: Flow cytometry gating strategy to analyze CD8+ T cell phenotype, related to Figure 5G-H. C. Soluble N-glycolylneuraminic acid and N-acetylneuraminic acid do not reduce Influenza A virus infection in Calu-3 lung epithelial cells, showing n=2 independent experiments, each an average of 3 wells. Baloxavir (influenza virus endonuclease inhibitor) is a positive control.

